# Comparative analysis of rhesus macaque and human placental organoids highlights evolutionary differences in placentation

**DOI:** 10.1101/2024.10.11.617873

**Authors:** Allyson Caldwell, Liheng Yang, Elizabeth A. Scheef, Amitinder Kaur, Carolyn B. Coyne

## Abstract

Throughout evolution, the placenta has diversified in structure and cellular composition while maintaining its essential role in supporting fetal development. Trophoblasts, key cells responsible for nutrient exchange and immune modulation, are a conserved feature of all placentas. Although primate placentas share broad morphological similarities, species-specific differences in gene expression remain poorly characterized, largely due to the lack of tractable *in vitro* models. To address this gap, we developed rhesus macaque placental organoids representing both trophoblast and maternal-derived decidual cell types and compared them with human placental organoids. Using integrated single-cell and single-nucleus RNA sequencing, we identified both shared and species-specific transcriptional programs across corresponding trophoblast lineages. We further reconstructed lineage trajectories leading to multinucleated syncytiotrophoblast and invasive extravillous trophoblast populations, revealing conserved differentiation pathways alongside divergent gene expression signatures. This work establishes new *in vitro* models of the nonhuman primate placenta and defines the molecular distinctions between human and rhesus trophoblasts, offering insights into the evolutionary adaptations underlying placental development.

## Introduction

The placenta forms the critical interface between mother and fetus during pregnancy, providing both nourishment and protection to the developing fetus. In hemochorial placentas, fetal trophoblast cells are in direct contact with maternal blood. Trophoblasts are essential for proper placental development and function, playing key roles in nutrient exchange, hormone production, and immune modulation. In hemochorial mammals such as humans and rhesus macaques, the trophoblast-derived chorion fuses with the allantois, an extra-embryonic membrane originating from the embryonic hindgut, to form the chorioallantoic placenta. As the chorioallantoic placenta has evolved, trophoblasts have been conserved over time^1^ and now include a variety of trophoblast populations. In most hemochorial mammals, these populations include cytotrophoblasts (CTBs) that can differentiate into other trophoblast subtypes including extravillous trophoblasts (EVTs) and the syncytiotrophoblast (STB). EVTs are located at the villous columns and invade into the decidualized endometrium where the placenta is attached during pregnancy. In early stages of gestation, EVTs remodel the maternal microvasculature to facilitate the entry of maternal blood into the intervillous space. The STB is the multinucleated trophoblast monolayer covering the surface of the hemochorial placenta and is formed by the fusion of mononuclear CTBs. While the overall architecture and trophoblast subtypes are conserved, key differences exist between species. For example, the extent and pattern of EVT invasion into the maternal decidua differs between humans and rhesus macaques^2^. These species-specific features underscore the importance of comparative models to study the maternal-fetal interface. The cellular heterogeneity and complexity of the placenta present a major challenge to accurately recapitulate placental biology *in vitro*, especially across species.

Rhesus macaques and humans develop morphologically similar placentas that contain comparable cell layers ^3^. These similarities, along with the close evolutionary relationship between rhesus macaques and humans, make rhesus macaques an excellent research model to study human placentation and pregnancy. However, despite their similarities, previous studies have identified key differences in the transcriptional signatures between rhesus macaque and human trophoblasts^4^. For example, while human trophoblast populations express markers including HLA-G, SIGLEC-6, and ERVW-1, these markers are absent in the rhesus macaque placenta^5–8^. Notably, these previous comparative studies were based on whole placental tissue rather than in distinct trophoblast cell populations.

Organoids are three-dimensional models that recapitulate key features of their tissue of origin, including cellular heterogeneity, organization, and function. Human trophoblast organoids have been validated as a model that accurately recapitulates the cellular heterogeneity within the human placenta and can be propagated and cryopreserved^9–11^. Our laboratory has previously developed human trophoblast organoids (hTO) and decidua organoids (hDO) from full-term placental tissue^11^. These organoids release pregnancy-associated hormones, differentiate to contain key trophoblast subpopulations present in the human placenta, and accurately recapitulate the transcriptional profile of trophoblasts observed *in vivo*^11,12^. Moreover, despite their isolation from full-term tissue, hTOs more closely resemble trophoblasts in first-trimester tissue^11,12^. These qualities make placental organoids a more physiologically relevant model than other *in vitro* models, such as cell lines derived from choriocarcinomas. In addition to tissue-derived organoids, trophoblast stem cells (TSC) have been developed as an *in vitro* model of both human and rhesus macaque placentas^13,14^. TSC models consist of proliferative mononuclear cytotrophoblasts that can differentiate to contain STB and EVT populations. While recapitulating aspects of trophoblast biology, there are differences between tissue-derived TOs and TSCs, with TSC-derived models lacking transcriptional signatures associated with trophoblast lineages^12,15,16^. Here, we developed rhesus macaque trophoblast organoids (rTO) and decidua organoids (rDO) from matched late gestation placentas. Morphologically, rTOs and rDOs closely resembled organoids derived from human tissue and expressed canonical markers of placental tissue. Moreover, these organoids recapitulated the hormone release and transcriptional signature of rhesus macaque placental tissue. We characterized the cellular heterogeneity of rTOs using single-cell (sc) and single-nucleus (sn) RNA sequencing and compared their transcriptional profiles to those of hTOs. Using this approach, we defined the transcriptional signatures of distinct rhesus macaque trophoblast populations and identified shared and unique gene signatures in human trophoblasts. To model differentiation, we treated rTOs and hTOs with media conditions known to promote EVT formation. We then compared the trajectories of cytotrophoblast (CTB) differentiation into STB and EVT lineages across species, revealing conserved and divergent transcriptional programs associated within these trophoblast subtypes. Altogether, this work establishes new *in vitro* models of the rhesus macaque placenta, allowing for direct cross-species comparisons to model human and non-human primate placentation.

## Results

### Derivation of trophoblast and decidua organoids from rhesus macaque placentas

To generate matched trophoblast and decidua organoids, we dissected fetal and maternal tissues from mid-late third trimester gestation rhesus macaque placentas (143-154 gestational days) (**Figure 1A**). The architecture of chorionic villi is similar between rhesus macaque and human placentas, but rhesus macaques exhibit shorter, wider, and less branched villi than do humans (**Figure 1B**). To develop rhesus macaque trophoblast organoids (rTOs), we dissected the chorionic villi and smooth chorion and carefully removed any decidua tissue. The chorionic villi and smooth chorion were homogenized using similar protocols that were developed previously to isolate human trophoblast organoids (hTOs) from full-term placental tissue^11^. To develop rhesus decidua organoids (rDOs), we isolated progenitor cells from decidual glands, following a similar approach as described for human DOs (hDOs). Isolated progenitor cells from the fetal and maternal tissues were then cultured in Matrigel to develop rhesus macaque placental organoids. Rhesus-specific growth factors are not commercial available therefore, rhesus macaque placental organoids were grown in previously defined term trophoblast organoid media (TOM) used for hTOs. Organoids were visible ∼5-7 days post-isolation and were passaged every ∼7-10 days (rTOs) or every ∼5-7 days (rDOs) once established. Morphologically, rTOs and rDOs resembled hTOs and hDOs (**Figure 1C**) and could be propagated long-term (>15 passages) without loss of this morphology (**Figure 1D**). Confocal microscopy for cytokeratin-19 confirmed the architecture of rhesus macaque organoids, which were similar to those derived from human placental tissue **(Figure S1A-B, D)**. Collectively, these data demonstrate the successful derivation of rTOs and rDOs from rhesus macaque placental tissue.

**Figure 1.**
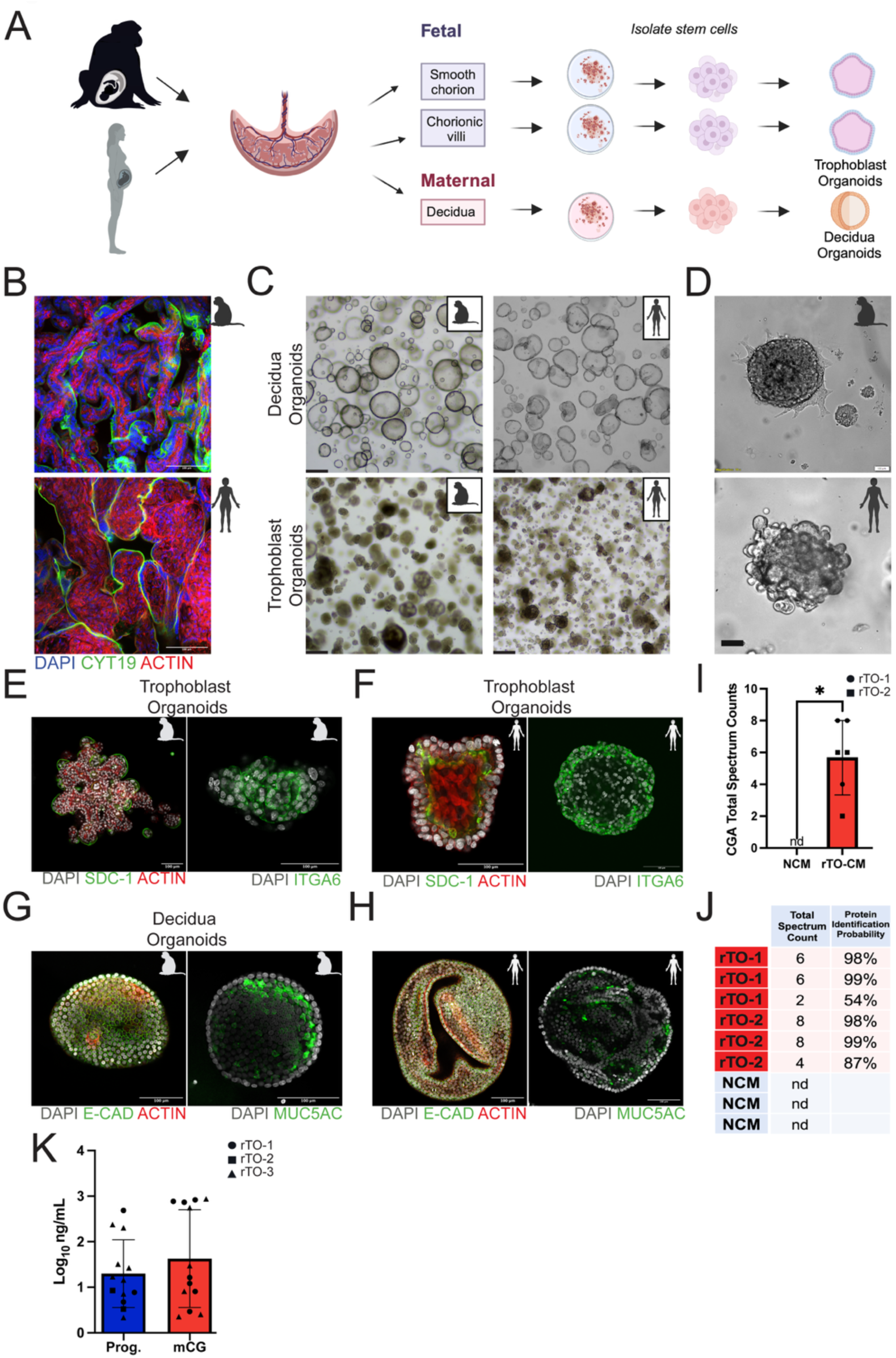
Establishment and of rhesus macaque trophoblast and decidua organoids. **(A),** Schematic of the rhesus macaque and human placenta denoting full-term human and mid-late third trimester rhesus macaque fetal trophoblast and maternal decidua tissues used to generate matched organoids. Created using Biorender. **(B),** Confocal micrographs of rhesus macaque (top) and human (bottom) chorionic villi. Cytokeratin-19 is shown in green, actin is shown in red, and DAPI-stained nuclei are in blue. Scale bar, 100μm. **(C),** Top, brightfield images of representative decidua organoids of rhesus macaque (left) and human (right). Bottom, representative brightfield images of trophoblast organoids of rhesus macaque (left) and human (right) trophoblast organoids. Scale bar, 500μm **(D),** Representative brightfield images of rTOs (top) and hTOs (bottom). Scale bar, 100μm. **(E, F),** Confocal micrographs of rTOs and hTOs immunostained for SDC1 (green), ACTIN (red), DAPI-stained nuclei are in black and white on the left, ITGA6 (green), DAPI-stained nuclei are in black and white on the right. Scale bar, 100μm. **(G, H),** Confocal micrographs of rDOs and hDOs immunostained for E-cadherin (green), ACTIN (red), DAPI-stained nuclei are in black and white on the left, MUC5AC (green), DAPI-stained nuclei are in black and white on the right. Scale bar, 100μm. **(I),** Total Spectrum Counts of Chorionic Gonadotropin (CGA) in non-conditioned media (NCM) and CM isolated from two independent lines of rTOs. Each point represents an independent replicate/well. **(J),** Quantification of total spectrum count and protein identification probability of mCG. **(K),** Secretion of progesterone and mCG from three unique rTO lines measured by radioimmunoassay. Each point represents an independent replicate/well.

### Rhesus macaque-derived placenta organoids secrete pregnancy hormones and express canonical markers

To examine the localization of canonical trophoblast and decidua markers in rTOs and rDOs, we performed immunostaining using cell-type specific markers. In rTOs, this included the STB marker syndecan-1 (SDC-1) and the CTB marker integrin alpha-6 (ITGA6). rTOs and hTOs both expressed SDC-1, indicating the presence of the fused STB and ITGA6 representing the mononuclear CTBs **(Figure 1E and 1F, S1C).** Unlike hTOs, which develop with an inward facing STB, SDC-1 localization suggested that rTOs developed with an outward facing STB **(Figure 1E and 1F**, **Supplemental Movies 1 and 2)**. Additionally, rDOs expressed decidua gland-associated markers including E-cadherin and the mucin MUC5AC, similar to their expression in hTOs **(Figure 1G and 1H, S1C)**.

Human TOs secrete essential pregnancy hormones including human chorionic gonadotropin (hCG)^9,11^. During pregnancy, the rhesus macaque placenta secretes chorionic gonadotropin (mCG) early in pregnancy with a peak occurring at approximately gestation day 22-24^17^. In addition, progesterone is secreted during rhesus macaque pregnancy similar to human pregnancy, with fluctuations throughout gestation^17^. Therefore, we investigated whether rTOs secreted mCG and/or progesterone. To do this, we used two approaches including unbiased mass spectrometry of conditioned medium (CM) collected from rTOs and an established radioimmunoassay^18,19^. For mass spectrometry, we collected CM from three independent wells of rTOs derived from two unique placentas and isolated bands migrating at ∼26kD and ∼10kD, the approximate size of chorionic gonadotropin A (CGA), on a reducing SDS-PAGE gel (**Figure S1E**). Spectral counts for CGA, which is secreted with CG beta subunits to form mCG, were detected in every conditioned media (CM) sample isolated from rhesus trophoblast organoids (rTOs), with high identification probability (**Figure 1I, 1J**). In contrast, no spectral counts for CGA mapped in any non-conditioned medium (NCM) control samples (**Figure 1I, 1J**). To directly quantify the levels of mCG and progesterone secreted from rTOs, we collected CM from three unique rTOs lines and performed a radioimmunoassay described previously^18,19^. NCM controls were assayed in triplicate and subtracted from rTO conditioned media samples. We found high levels of mCG (>1000ng/mL) and progesterone (>1200ng/mL) in rTO conditioned media, **(Figure 1K)**. Similar to hTOs, rTOs can be passaged long-term without loss of mCG or progesterone secretion.

Lastly, we performed bulk RNA sequencing of rTOs and rDOs to assess the expression of known canonical markers of human trophoblasts or glandular epithelial cells, respectively, and to verify the absence of contamination of other tissues/cells. To do this, we compared the expression profile of canonical markers of rhesus-derived trophoblasts and decidua epithelial organoids. We found that rTOs highly expressed trophoblast-specific markers, including *PGF*, *MMP2*, and *ITGA5* which were absent or expressed at low levels in rDOs **(Figure S1F)**. We also found that rTOs expressed markers of distinct trophoblast lineages including the STB (e.g., *CGA*), CTBs (e.g., *FOXO4*), and EVTs (e.g., *MAMU-AG*). Conversely, rDOs expressed markers associated with the maternal decidua including *MUC1*, *MUC20*, and *SOX17*, which were absent or expressed at low levels in rTOs **(Figure S1F, S1J)**. These results were similar to the differences observed between hTOs and hDOs **(Figure S1G)**.

We next analyzed the transcriptional dataset to identify non-annotated genes associated with rhesus macaque trophoblasts, which included pregnancy-specific glycoproteins (PSGs). PSGs have been described in many non-human species, including primates, mice, horses, and cows^20^; however, the driving forces of evolution of these genes is unclear. Rhesus macaque PSGs have not been annotated with gene names in the recent *Macaca Mulatta* genome assembly ( Mmul10). We identified rhesus macaque PSGs in the rTO dataset by comparing the genome sequence of unannotated macaque genes to the human genome (hg38) using BLAST. We identified rhesus macaque PSGs in the rTO dataset by comparing the genome sequences of unannotated macaque genes to the human reference genome (hg38) using BLAST. This analysis revealed five PSGs that were expressed in rTOs (**Figure S1H**). In comparison, ten PSGs were expressed in hTOs based on annotations in the human genome (**Figure S1I**). Together, these data demonstrate that rTOs and rDOs recapitulate key aspects of the rhesus macaque placental transcriptome, including the expression of cell type–specific markers.

To characterize the transcriptional profiles of rhesus TOs and DOs, we performed differential gene expression analysis using edgeR to identify genes distinguishing these organoid types (**Figure S1J**). This analysis revealed distinct expression profiles between rTOs and rDOs. To confirm the identity of decidual organoids, we also conducted signature scoring using a curated set of glandular epithelial markers, which demonstrated robust expression of decidual-associated genes in rDOs (**Figure S1K**). We next compared gene expression profiles between rDOs and hDOs using edgeR to identify both species-specific and conserved transcriptional signatures (**Figure S1L**). This analysis revealed clear divergence in gene expression patterns between species. Notably, *TRIM72* and *PARP6* were significantly upregulated in rDOs, while *OST4* and *EMC4* showed higher expression in hDOs, underscoring species-specific molecular programs within decidual tissues. Additional differentially expressed genes included cytokines, mitochondrial components, and transcriptional regulators, reflecting both functional specialization and evolutionary divergence. To investigate the functional relevance of genes conserved between rDOs and hDOs, we performed gene ontology (GO) enrichment analysis using the clusterProfiler algorithm (**Figure S1M**). This analysis revealed significant enrichment in pathways related to olfactory receptor activity, DNA-binding transcription activator activity, cytokine activity, and receptor ligand activity. These findings emphasize both the evolutionary conservation and divergence of decidual biology across primates and support the utility of the rDO model for cross-species comparisons.

### Single-cell RNA-seq mapping of rTOs reveals comparable cellular heterogeneity to hTOs

To evaluate whether rTOs mirror the cellular heterogeneity of the rhesus placenta, we performed single-cell RNA sequencing (scRNAseq) on rTOs cultured under standard conditions. In parallel, we generated new scRNAseq data from hTOs and incorporated previously published datasets from our prior work^12^. Established lines of rTOs and hTOs were dissociated into single cell suspensions and processed separately to identify cell types present in TOs. A total of 11,916 (rTOs) and 15,766 (hTOs) cells passed quality control and were used for cluster analysis and cell-type specific marker expression analysis following integration to correct for batch effects. Clustering analysis revealed five major cell populations present in both rTOs and hTOs (**Figure 2A, 2B**) which were distributed similarly across all TO lines (**Figure 2C, 2D, S2A, S2B**). To define the cellular identity of each cluster, we identified cluster-enriched gene expression profiles and mapped them to canonical trophoblast markers^4,14^. Clusters in rTOs included two non-proliferative CTB populations (CTB-1 and -2) which accounted for ∼31% and ∼20% of all populations respectively, two proliferating CTB populations (CTBp-1 and -2) which accounted for ∼24% and ∼23%, and one STB cluster which only accounted for 2% (**Figure 2A, 2C**). Given the multinucleated nature of the STB, it is less represented in scRNAseq datasets from both tissue and TOs^12^. CTB clusters were enriched in canonical markers including *KRT7* and *CDH1* whereas CTB-p clusters were enriched in genes associated with proliferation (e.g., *MKI67*, *CCNA2*) (**Figure 2E 2G**). The STB cluster was enriched in canonical markers including *TFAP2A*, *INSL4*, *HOPX*, *CGA*, and the endogenous retrovirus fusion gene *ERVFRD-1* (**Figure 2E, 2G**). Many genes previously identified in TSC-derived rhesus macaque lineages^14^ to be expressed in CTBs (**Figure S2C**) and STB/EVTs (**Figure S2D**) were also expressed in a lineage-specific manner. Unbiased cluster gene enrichment analysis also demonstrated that as expected, both CTB-p clusters were enriched for genes associated with cell division (e.g., *CENPF*, *CCNB1*) and histones (e.g., *H1-5*, *H1-3*) (**Figure S2E**). In contrast, the CTB-1 cluster in rTOs was enriched in gene associated with extracellular matrix (e.g., *LAMB3*, *LAMC2*) and the regulation of MMPs (e.g., *TIMP1*, *TIMP3*) whereas CTB-2 was enriched in genes associated with GTPases and GTP-binding proteins (e.g., *ARL15*, *AGAP1*) and NOTCH signaling (e.g., *MAML2*, *MAML3*) (**Figure S2E**). These expression profiles were distinct from genes enriched in the STB, which included canonical markers (e.g., *HOPX*, *CGA*) and several non-annotated genes (e.g., *ENSMMUG00000049714*, *ENSMMUG00000063643*) (**Figure S2E**). Many of the non-annotated STB-enriched genes are predicted to encode for rhesus macaque CGBs (*ENSMMUG00000040968, ENSMMUG00000063643, ENSMMUG00000049714*) and PSGs as we described above (*ENSMMUG00000039210, ENSMMUG00000050799, ENSMMUG00000047533*) (**Figure S2F, S2G**). Previous studies described the expression of CGBs in macaque trophoblast stem cell lines^14,21^ but these studies did not state which specific CGBs were present. The GO terms and genes associated with population-specific trophoblast-enriched genes included the regulation of cell locomotion, motility, and migration, actin cytoskeletal organization and processes (e.g., *ACTG1*, *PFN1*, *CDC42*) in CTB-1 whereas those associated with CTB-2 were involved in the regulation of GTPase and hydrolase activities (e.g., *RGL1*, *SNX13*, *RALGPS2*, *ARFGEF1*) (**Figure S2H, S2I**). These pathways and genes were distinct from the STB, in which pathways including protein catabolic processes and endomembrane system organization were enriched (e.g., *FOLRF1*, *EHD3*, *RAB11A*) (**Figure S2H, S2I**). STB-associated GO terms and associated genes included catabolic processes, synaptic vesicle maturation, and protein localization to the plasma membrane (**Figure S2H, S2I**). These data highlight the distinct cellular processes and functions amongst distinct rhesus macaque trophoblast populations in rTOs.

**Figure 2.**
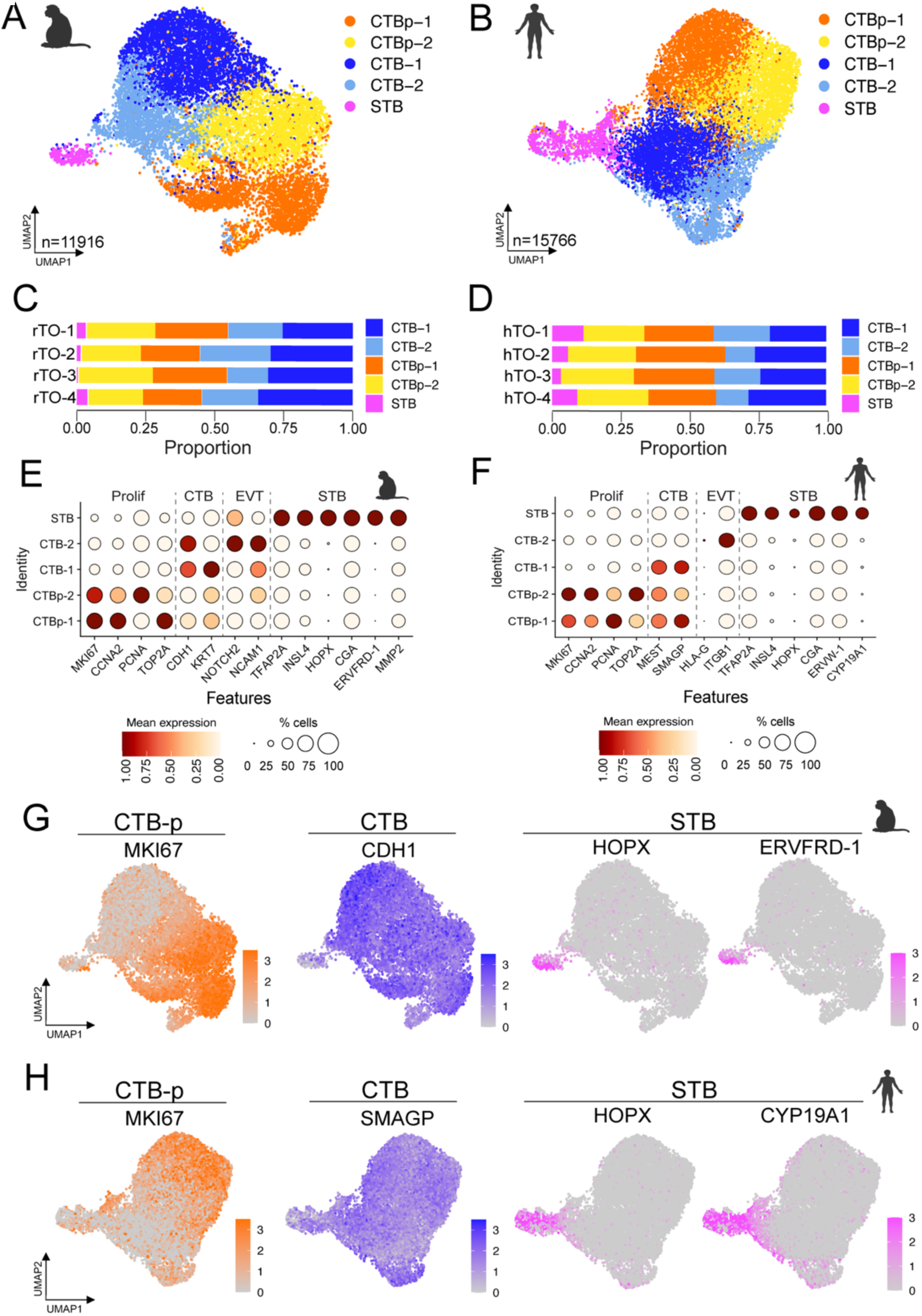
Single cell RNA-seq defines trophoblast cell populations present in rhesus macaque and human trophoblast organoids. **(A, B),** UMAP of cell type clusters from four lines of rhesus macaque (rTOs, A, n=11,916 cells) and four lines of human (hTOs, D, n=15,776 cells) trophoblast organoids. Single-cell dataset generated from three unique lines of rTOs and four unique lines of hTOs. Cell types are denoted as follows: cytotrophoblasts (CTB), proliferative cytotrophoblasts (CTB-p), extravillous trophoblasts (EVT), syncytiotrophoblasts (STB). Multiple clusters of one cell type are annotated using -number after the cell name. **(C, D),** Comparison of the enrichment of distinct cell clusters in rTOs (C) and hTOs (D). **(E, F),** Dot plots showing expression levels of indicated genes in each cluster in rTOs (E) and hTOs (F). The indicated genes are established markers for the syncytiotrophoblast (STB), cytotrophoblasts (CTBs), extravillous trophoblasts (EVTs) and trophoblast progenitor cells (CTB-p). Scale is shown at bottom. **(G, H),** Feature plots of trophoblast marker gene expression in rTOs (G) or hTOs (H). Scale is shown at right.

Clusters in hTOs included two non-proliferative CTB populations (CTB-1 and -2) which accounted for ∼24% and ∼15% of all populations respectively, two proliferating CTB populations (CTBp-1 and -2) which accounted for ∼27% and ∼24%, and one STB cluster which only accounted for 8% (**Figure 2B, 2D**). Gene enrichment in hTOs was similar to rTOs, with pCTBs highly enriched for markers of proliferation (e.g., *MKI67* and *CCNA2*), CTBs enriched for canonical markers (e.g., *KRT7* and *CDH1*), and the STB enriched for canonical marker (e.g., *TFAP2A, INSL4, HOPX, ERVW-1*, and *CYP19A1*) **(Figure 2F, 2H)**. Also, like rTOs, one CTB cluster, CTB-2, expressed markers of undifferentiated EVTs including *ITGB1* and expressed low levels of the canonical EVT marker *HLA-G* (**Figure 2F**). Collectively, these data define the cellular composition of rhesus macaque- and human derived-trophoblast organoids and show that these organoids differentiate into multiple trophoblast lineages.

### Defining trophoblast marker divergence and convergence in rTOs and hTOs

Next, we integrated scRNAseq datasets from rTOs and hTOs to perform evolutionary gene expression analysis, aiming to identify both divergence and convergence in cell marker expression between the two species. This analysis revealed five conserved clusters, most of which were comprised of near-equivalent ratios of rTO- and hTO-derived cells (**Figure 3A, 3B**). These populations were conserved across independent rTO and hTO lines (**Figure S3A**). Clusters expressed canonical markers of CTB-p (e.g., *MKI67*, *TOP2A*), CTBs (e.g., *CDH1*, *KRT7*), and the STB (e.g., *TFAP2A*, *HOPX*) (**Figure 3C**). Expression of these canonical markers was conserved in both rTO and hTO datasets (**Figure 3D**). Unbiased gene enrichment analysis identified select genes defining two CTBp clusters (CTBp-1, CTBp-2), two CTB clusters (CTB-1, CTB-2) and a single STB cluster (**Figure S3B**). We did not observe distinct EVT clusters in the combined dataset but select clusters did express both rhesus macaque and human EVT markers **(Figure S3C)**. We performed gene ontology enrichment analysis, using the *clusterProfiler* algorithm, on rhesus macaque and human datasets and identified molecular function pathways uniquely enriched in each (**Figure S3D, 3E, 3F)**. Next, we investigated the transcriptional profiles of the CTBp and CTB clusters (**Figure S3G–J**). Notably, we found that CTBp clusters exhibited a higher degree of transcriptional conservation across species, with a greater proportion of shared marker genes compared to CTB clusters. This suggests that the proliferative state of cytotrophoblasts may be governed by more conserved regulatory programs, whereas non-proliferative CTBs may exhibit greater species-specific variation in gene expression.

**Figure 3.**
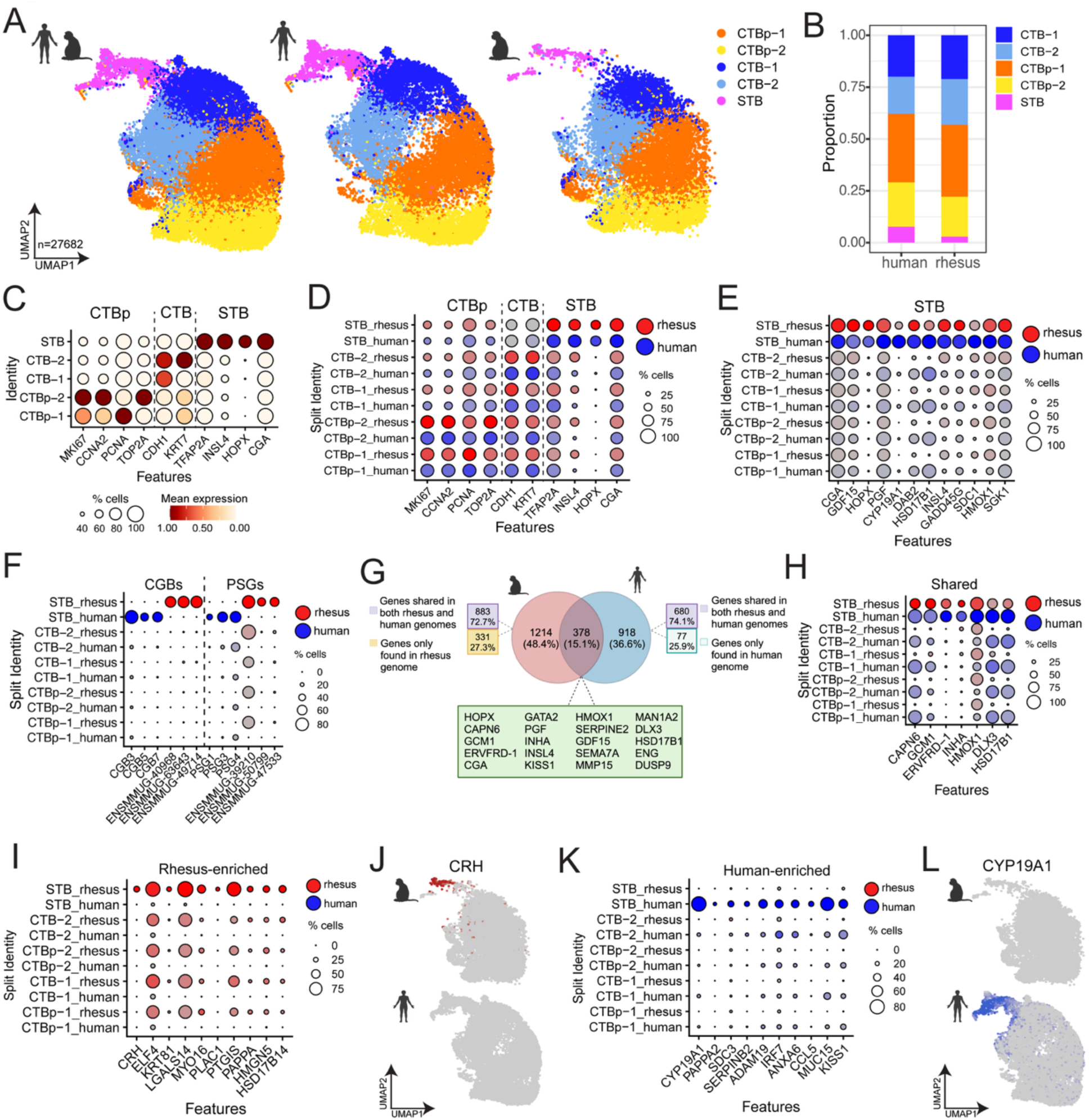
Integrated analysis of rhesus macaque and human single cell RNA-seq identifies shared and distinct cellular transcriptional profiles. **(A),** UMAP of merged cell type clusters from integrated datasets rTOs and hTOs (n=27,682 total cells). Three unique lines of rTOs and four unique lines of hTOs were used with four total samples from each species. Left, merged UMAP of clusters from rTOs and hTOs. Middle, cells from hTOs and right, cells from rTOs. **(B),** Cluster enrichment from hTO (left) and rTO (right) present in the integrated dataset. **(C),** Dot plots of canonical trophoblast markers in each cluster present in the integrated dataset. Key at bottom. **(D, E),** Canonical trophoblast markers in each cluster (D) or in the STB (E) present in the integrated dataset separated by species with rhesus in red and human in blue. Key at right. **(F),** Dot plots of rhesus macaque and human CGBs and PSGs (rhesus in red and human in blue). Key at right. **(G),** Venn diagram of markers in the rhesus and human STB clusters that identifies shared and distinct markers. Markers that were shared are shown as genes that are present in both the rhesus macaque and human genome. Distinct rhesus and human STB markers include genes that are shared in the rhesus macaque and human genome as well as genes that are specific to one genome. **(H, I, J, K, L),** Dot plots of shared (H), rhesus-enriched (I) or human-enriched (K) markers found in the STB cluster separated by rTO and hTO conditions (rhesus in red and human in blue). Feature plot of CRH (J) and CYP19A1 (L) separated by rTO and hTO conditions.

We next focused on differential gene expression within the STB cluster between rTOs and hTOs. While this cell type is conserved across species, the extent to which its transcriptional profile diverges remains largely unknown. We found that there was near-equivalent expression of canonical STB genes including *HOPX*, *INSL4*, *CGA*, in both rTOs and hTOs (**Figure 3E**). We also observed the expression of orthologs in the STB cluster, including genes such as CGBs and PSGs that were expressed at similar levels in both rTOs and hTOs (**Figure 3F**). To elucidate differences in the transcriptional profiles of the STB markers rTOs and hTOs, we performed pseudobulk differential expression analysis using DESeq2. This analysis identified differences in STB-associated transcriptional profiles and highlighted variations in gene expression arising from differences in annotation and/or gene presence between the human and rhesus genomes **(Figure 3G)**. Of the STB markers in the integrated dataset, 15.1% were shared between the two species **(Figure 3G)**. These markers included calpain-6 (*CAPN6*), inhibin subunit alpha (I*NHA*), and *ERVFRD-1* (**Figure 3H**). rTO-specific STB markers accounted for 48.4% of the STB markers and 27.3% of these genes were found only in the rhesus macaque genome (**Figure 3G**). hTO-specific STB markers made up 36.6% of the STB markers and 25.9% of these markers were found only in the human genome (**Figure 3G**). We therefore focused on defining gene expression conservation and variance from genes conserved in both genomes. This analysis revealed significant differences in STB marker expression between rTOs and hTOs. For example, the STB in rTOs expressed the peptide hormone corticotropin-releasing hormone (*CRH*), the transcription factor *ELF4*, and the cytokeratin *KRT81*, amongst others (**Figure 3I, 3J**). In contrast, hTOs were enriched in genes including *CYP19A1*, the endogenous retrovirus gene *ERVV-1*, and the chemokine *CCL5* (**Figure 3K, 3L**). These data demonstrate that although rTOs and hTOs share key markers within their STB cells, each also exhibits species-specific gene expression signatures within this specialized cell type.

### Single-nucleus RNA Sequencing Expands Resolution of Trophoblast Cell States in rTOs

Using scRNAseq, we identified both shared and species-specific features of STB cells in rTOs and hTOs. However, prior studies have shown that snRNAseq is essential for accurately capturing gene expression in the STB, due to their multinucleated nature and the limitations of scRNAseq in profiling such cells^12,16^. Therefore, to further characterize the specialized trophoblast populations in rTOs and their transcriptional profiles, we performed snRNAseq on rTOs. Three lines of rTOs were dissociated into single cell suspensions and nuclei isolated. To compare trophoblast populations captured by the two sequencing modalities, we integrated rTO scRNAseq and snRNAseq datasets (**Figure 4A**). This integrated dataset revealed that, consistent with previous studies in hTOs^12^, scRNAseq of rTOs captured fewer STB-associated cells than snRNAseq (**Figure 4B, Figure S4A**). STB-associated cells comprised only 1.92% of the scRNAseq dataset compared to 15.3% of the snRNAseq dataset, underscoring the enhanced detection of this multinucleated population by nuclear profiling (**Figure 4B**). UMAP projection of the combined dataset identified eight trophoblast clusters, including two proliferating CTB (CTBp) clusters, five non-proliferating CTB clusters, and one STB cluster, defined based on canonical subtype markers (**Figure S4B**). Proliferative CTBs were overrepresented in s-RNAseq, accounting for 41.6% of cells versus 27.3% in snRNAseq. Despite these modality-specific biases, non-proliferating CTBs remained the dominant population in both datasets, representing 56.5% and 57.4% of cells in sc-RNAseq and snRNAseq, respectively (**Figure 4B**).

**Figure 4.**
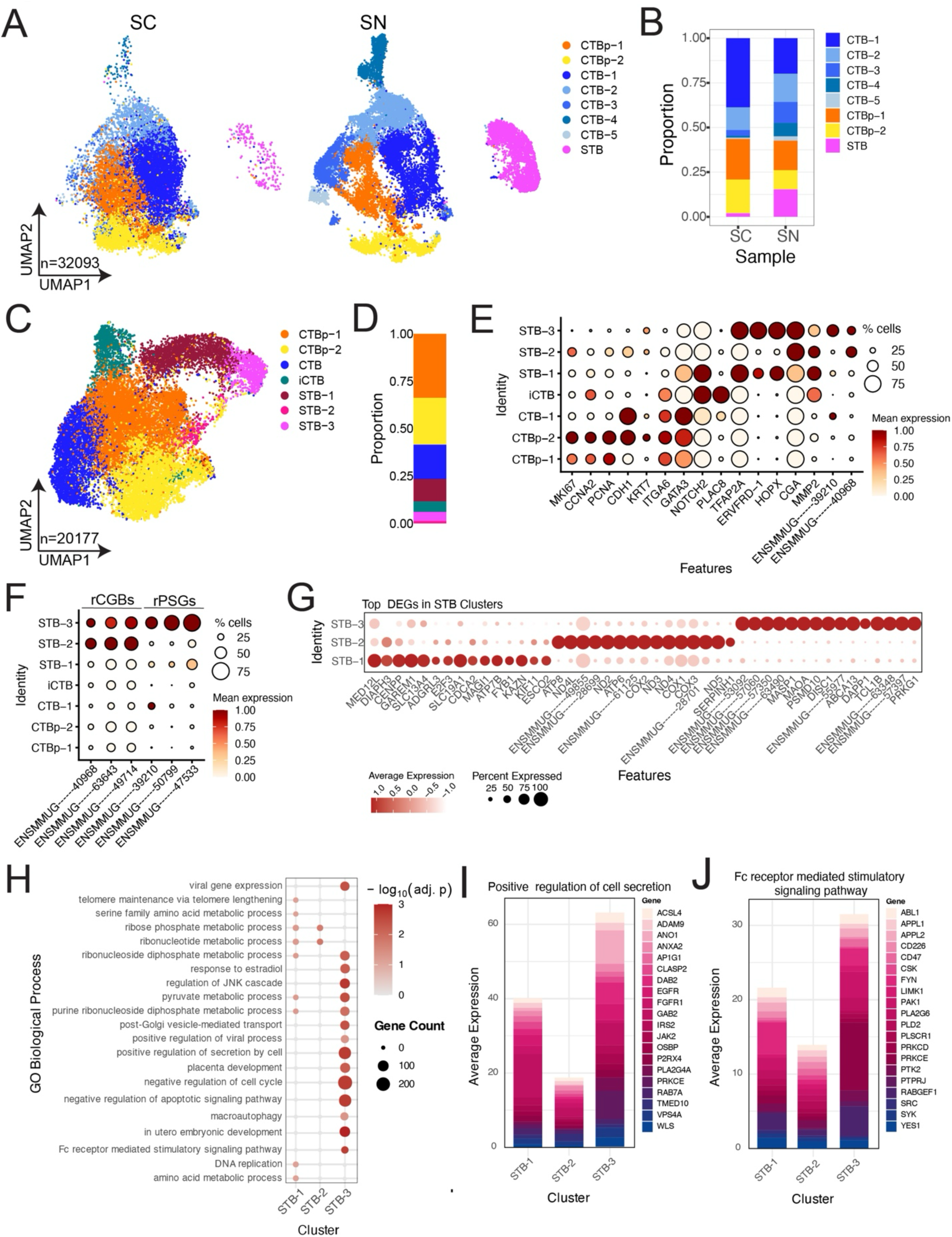
Single-nucleus RNA Sequencing captures the full diversity of trophoblast cellular and transcriptional profile of rhesus macaque and human trophoblast organoids. **(A),** UMAP of integrated datasets of rTO single-cell (SC) and single-nucleus (SN) RNA sequencing (n= 32,093 total cells/nuclei). Single-cell dataset generated from four rTO samples (SC, left) and single-nucleus generated from three rTO samples (SN, right). **(B),** Enrichment of cell clusters in SC (left) and SN (right). **(C),** UMAP of rTO SN dataset (n= 20,177 nuclei) created from three unique rTO lines. Seven clusters were identified, and cell types were indicated as follows: proliferative cytotrophoblasts (CTB-p), cytotrophoblasts (CTB), invasive cytotrophoblasts (iCTB), syncytiotrophoblast (STB). **(D),** Enrichment of cell clusters in rTO SN dataset depicted as a bar plot. **(E),** Dot plot depicting the expression levels of trophoblast markers shown in each cluster. Key at right. **(F),** Dot plot of rhesus CGBs and PSGs expression in each cluster. Key at right. **(G),** Dot plot of top differentially expressed genes in STB clusters. Key at bottom. **(H),** Dot plot of pathways from GO enrichment in STB clusters. Key at right. **(I, J),** Bar plots illustrating average expression of genes associated with positive regulation of cell secretion pathway (I) and Fc receptor mediated stimulatory signaling pathway in STB clusters.

Given the improved detection of the STB by snRNAseq, we focused our subsequent analysis on the snRNAseq dataset, as our prior work demonstrated that this sequencing modality more effectively resolves STB heterogeneity in hTOs^12^. We sought to determine whether a similar degree of complexity was present in rTOs. After quality control filtering, 20,177 rTO nuclei were retained for analysis. Clustering revealed seven distinct trophoblast populations: CTBp clusters, one non-proliferative CTB cluster, one invasive CTB (iCTB) cluster, and three STB clusters (**Figure 4C, 4D, S4C**). These clusters expressed canonical markers of distinct trophoblasts, including markers of proliferation (e.g., *MKI67*, *CCNA2*), CTBs (e.g., *CDH1*, *KRT7*), invasive CTBs (e.g., *ITGA6*, *NOTCH2*), and STB (e.g., *HOPX*, *CGA*, *ENSMMUG00000039210*) (**Figure 4E**). Cluster-specific expression patterns were also evident. For example, *COL15A1*, *LAMB4*, and *PTGER3* marked the iCTB cluster; *TAFA2*, and *TIMP3* were enriched in CTBp-1; while *HOPX*, *FYN*, and *ENSMMUG00000057397* were selectively expressed in various STB clusters (**Figure S4D**). These data highlight the transcriptional heterogeneity of rTO trophoblasts and support the presence of discrete subpopulations within the STB lineage.

To further investigate the differences among STB subtypes, we examined the expression of the RM *CGB* and *PSG* genes identified above. These genes were selectively expressed in the STB-2 and STB-3 clusters, suggesting that these populations represent more differentiated STB states (**Figure 4F**). In contrast, the STB-1 cluster exhibited high expression of *ERVFRD-1*, an endogenous retroviral gene known to mediate trophoblast cell fusion (**Figure 4E**). This pattern suggests that STB-1 may represent an immature STB cluster and/or a transitional state undergoing fusion from CTB to STB. To dissect transcriptional differences among STB subpopulations, we examined the top differentially expressed genes (DEGs) in each STB cluster. STB-2 and STB-3 showed strong expression of rhesus *CGBs* (*ENSMMUG00000040968*, *ENSMMUG00000063643*, *ENSMMUG00000049714*) and *PSGs* (*ENSMMUG00000039210*, *ENSMMUG00000050799*, *ENSMMUG00000047533*) (**Figure 4G**). In contrast, STB-1 lacked these markers and instead expressed *MED12L*, *DIAPH3*, and *CENPP*, genes associated with transcriptional regulation, cytoskeletal remodeling, and cell division. These features suggest that STB-1 represents an immature or transitional syncytial state. STB-3 was distinguished by expression of several uncharacterized macaque-specific transcripts (*e.g.*, *ENSMMUG00000039366*, *ENSMMUG00000039391*), highlighting potential species-specific features of terminal STB differentiation.

To define functional differences among STB subtypes, we performed GO enrichment analysis on cluster-specific differentially expressed genes. This analysis revealed distinct biological programs associated with each subtype (**Figure 4H**). STB-1 was strongly enriched for cell cycle–related pathways, including chromosome segregation, DNA replication, and mitotic cell cycle phase transition (e.g., *CENPP*, *CCNE1*, *WEE1*, *CHEK2*), consistent with a proliferative or fusion-competent syncytial precursor population. STB-2 was enriched for processes related to vesicle trafficking, including post-Golgi vesicle-mediated transport. Representative genes included *SEC22B*, *RAB27A*, *TMED10*, and *WLS*, indicating specialization in regulated secretion, possibly of hormones or signaling molecules. STB-3 was enriched for pregnancy- and immune-regulatory pathways such as Fc receptor–mediated stimulatory signaling, regulation of leukocyte activation, response to estradiol, placenta development, positive regulation of cell secretion, and in utero embryonic development (**Figure 4H**). The top 20 genes associated with two of these pathways, positive regulation of cell secretion and Fc receptor mediated stimulatory signaling pathway, show an enrichment of these genes specifically in STB-3 (**Figure 4I, 4J**). Together, these enrichment results support a model in which STB-1 represents a cycling or fusion-ready precursor population, STB-2 is specialized for vesicular sorting, and STB-3 carries out immune-modulatory and pregnancy-associated functions. This functional stratification underscores the transcriptional and biological heterogeneity of the STB within rTOs.

### Pseudotime Analysis Identifies Distinct Stages of STB Differentiation and Function in rTOs

To compare the progression of STB differentiation in rTOs and hTOs, we performed trajectory analysis using previously published hTO datasets^12^ alongside rTO data. Pseudotime analysis is a powerful approach for modeling gene expression dynamics along differentiation lineages and identifying regulatory genes and pathways that drive cell state transitions. We applied the Slingshot algorithm to infer pseudotime trajectories from sc- and sn-RNA-seq datasets, enabling reconstruction of the CTB to STB differentiation continuum in both species. Global lineage structures were inferred without predefining root or terminal clusters, allowing unbiased trajectory detection across trophoblast states. We visualized the resulting pseudotime trajectories on UMAP embeddings of rTO and hTO datasets derived from snRNAseq (**Figure 5A–B, S4E**) and scRNAseq (**Figure S5A–B**). In both modalities, we identified a continuous developmental trajectory originating from proliferative CTBs (CTB-p), passing through non-proliferative CTBs, and culminating in the STB cluster (**Figures 5A, 5B; S5A, S5B, S5C, S5E; Table S8-9**). In rTOs, the inferred lineage structure revealed two distinct branches emerging from the CTB compartment: one progressing from STB-1 directly to STB-3, and a second, independent branch terminating in STB-2 (**Figure 5A, Figure S4E**). This suggests that STB-2 may represent a distinct differentiation pathway or functional state, rather than a transitional intermediate. To further define gene expression dynamics along these trajectories, we used tradeSeq to model pseudotime-dependent expression changes in both rTOs and hTOs. In rTOs, canonical STB markers such as *GCM1* and *MAN1A2* increased along the STB-1 to STB-3 trajectory (**Figure 5C**), while expression of CTB markers including *CDH1* declined (***Figure 5G*, S5G**). *ERVFRD-1*, an endogenous retroviral fusion gene associated with trophoblast syncytialization, was also upregulated along this path (**Figure 5G**). In hTOs, a similar increase in STB markers (e.g., *CYP11A1*, *PSG5*, and *HOPX)* was observed along the pseudotime axis (**Figures 5D, 5F, S5D, S5F, Table S10-11**), accompanied by a loss of *CDH1* expression and an increase in *PSG9* (**Figure 5H, S5H**). Together, these data define conserved and divergent gene expression dynamics underlying CTB-to-STB differentiation in human and rhesus TOs. The presence of branching trajectories in rTOs suggests subtype-specific regulatory programs and highlights the power of pseudotime analysis to resolve complex lineage relationships during STB development.

**Figure 5.**
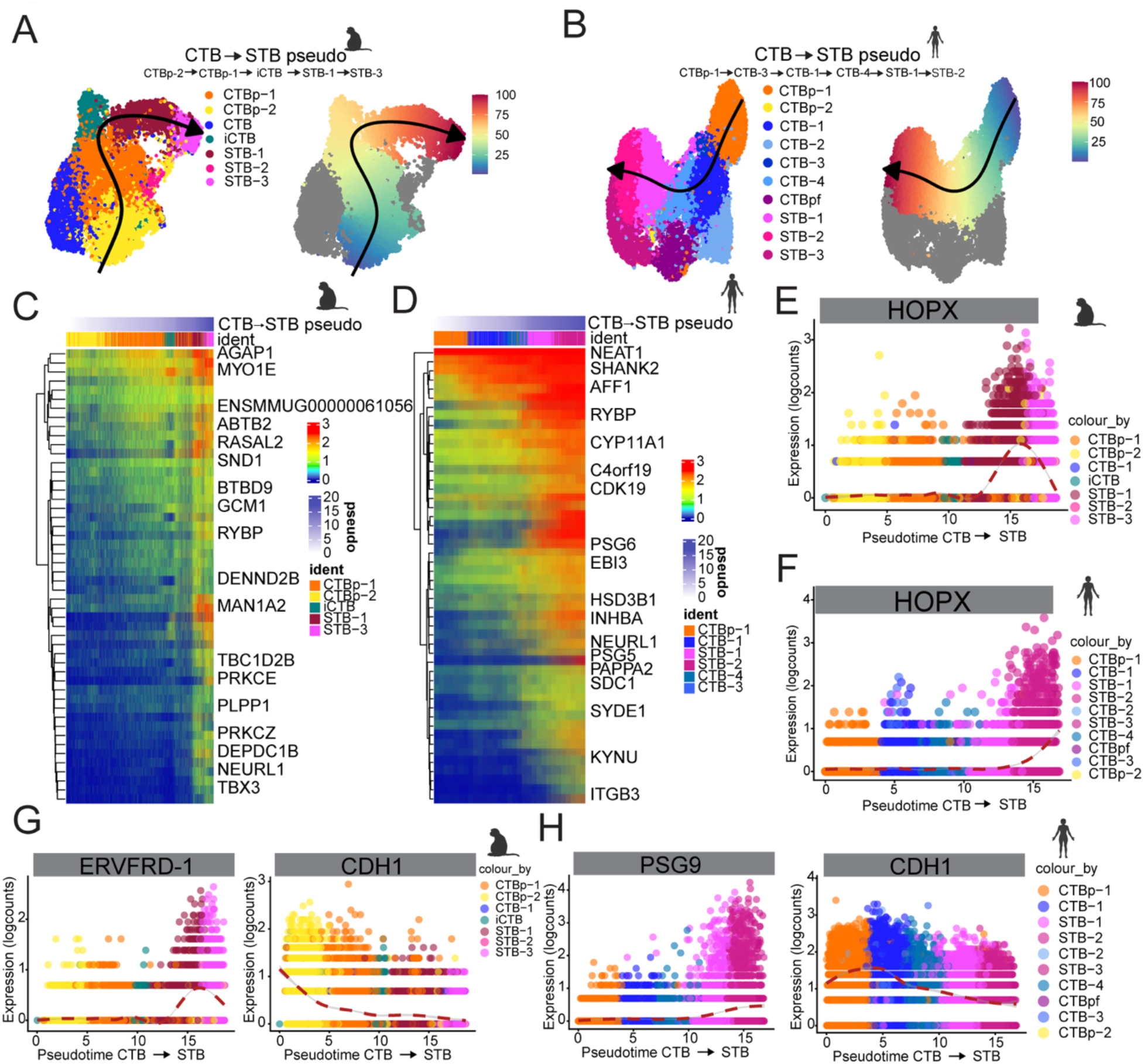
Inter-species profiling of rhesus macaque and human syncytiotrophoblast trajectory in sn-RNAseq datasets. **(A, B),** To analyze cellular lineage of trophoblast in rTOs and hTOs, pseudotime was implemented to visualize the trajectory of trophoblast cell types rTO (A, right) and hTO (B, right). Pseudotime values were assigned to each cell type and input into a color scale matching the UMAP based on time 0-100 in rTO (A, left) and hTO (B, left). The arrow across the UMAP shows the order of the trajectory. Pseudotime values revealed top genes correlated to the trajectory in rTO (A, right)and hTO (B, right). **(C, D),** Heatmap of the top increasing genes associated with the STB trajectory enriched in rTO (C) and hTO (D) dataset. **(E, F),** Increase of HOPX gene expression along the differentiation trajectory to STB subtype in rTO (E) and hTO (F). The cell types presented are represented by the colors denoted in the legend. Logcounts and pseudotime values are shown as a color scale present in the legend. **(G),** Normalized expression of ERVFRD-1and CDH1 shown along rTO pseudotime. **(H),** Normalized expression of PSG9 and CDH1 shown along hTO pseudotime.

### Pseudotime analysis uncovers species-specific gene trajectories in the STB lineage between rTOs and hTOs

Given the significant differences in gene expression profile of STB cells between rTOs and hTOs, we conducted a comparative Slingshot pseudotime trajectory analysis using the integrated rTO and hTO scRNAseq dataset to infer gene expression along the CTBp to STB differentiation. This pseudotime analysis revealed a continuous STB trajectory, progressing from proliferative CTB to non-proliferating CTB and finally to the differentiated STB subtype (**Figure S6A**). STB differentiation was characterized by increased expression of well-known STB markers *INSL4*, *PDLIM2*, *HMOX1*, *GCN1*, and *ERVFRD-1*, which were upregulated in both rTOs and hTOs **(Figure S6B, S6C, S6D, Table S12)**. Next, we compared species-specific gene trajectories between rTOs and hTOs by analyzing the integrated dataset, separating the data by condition. This analysis revealed species-associated differences in gene expression along the STB trajectory in rTOs and hTOs. The rTO gene trajectory showed increased expression of *PAGE4*, *SLC2A1*, *WNT5A*, and *NGF* in the STB trajectory, while these genes were unchanged or downregulated in the hTO trajectory (**Figure S6E-S6H**). Conversely, *SDC3*, *TGFB1*, *HSPB1*, and *ESAM* specifically increased only in the hTO STB trajectory, with no similar increase in rTOs (**Figure S6I-S6L**). Altogether, these findings highlight species-specific differences in gene expression dynamics along the STB differentiation trajectory.

### Transcriptional Profiling of EVT Differentiation in Rhesus Trophoblast Organoids

In rTO scRNAseq and snRNAseq datasets, a distinct EVT cluster was not detected, consistent with prior findings in hTOs under basal culture conditions^12^. To determine whether EVT differentiation could be induced in rTOs, protocols previously developed for hTOs^11,13,22^ were adapted and tested in the rhesus system. This multi-step differentiation process, which includes treatment with Neuregulin-1 (NRG1), has been shown to promote EVT fate in human trophoblast organoids^11,13,22^. Following induction, snRNAseq was performed on three independent rTO lines to evaluate the extent of EVT differentiation and associated transcriptional changes. To assess whether EVT differentiation was successfully induced, snRNAseq data from control (mock-treated) and EVT-differentiated (EVT^enrich^) rTOs were integrated for comparative analysis. The combined dataset revealed eight transcriptionally distinct clusters, including two proliferative CTB clusters (CTBp), three non-proliferative CTB clusters, one STB cluster, and one prominent EVT cluster based on canonical gene expression (**Figure 6A-6C**). Most CTBp and CTB populations were present at comparable frequencies between conditions. For example, CTBp-1 accounted for 27.8% of nuclei in EVT^enrich^ rTOs and 28.7% in controls, while CTBp-2 and CTBp-3 remained under 6% in both conditions (**Figure 6B**). Similarly, CTB-1 and CTB-2 were detected at roughly 10–13% in both groups. The primary difference within the CTB compartment was CTB-3, which was more abundant in mock-treated rTOs (24.4%) compared to EVT^enrich^ samples (17.0%) (**Figure 6B**). A marked expansion of the EVT cluster was observed in the EVT^enrich^ condition, where EVT cells made up 21.5% of total nuclei, compared to just 4.33% in mock-treated rTOs (**Figure 6A, 6B**). In contrast, the STB cluster was reduced following EVT differentiation, comprising 2.87% of nuclei in EVT^enrich^ rTOs compared to 13.5% in mock-treated controls, similar to previous findings in hTOs, where STB representation similarly declined under EVT-inducing conditions^12^. The EVT cluster was defined by expression of canonical rhesus EVT markers previously described *in vivo*^14^, including *TIMP2*, *NOTCH2*, *GRHL1*, and *NCAM1* (**Figure 6C**). While *MAMU-AG*, a rhesus-specific homolog of *HLA-G*, is a well-established EVT marker, it was not detected in our dataset, likely due to limitations in transcript annotation or insufficient read coverage in this region.

**Figure 6.**
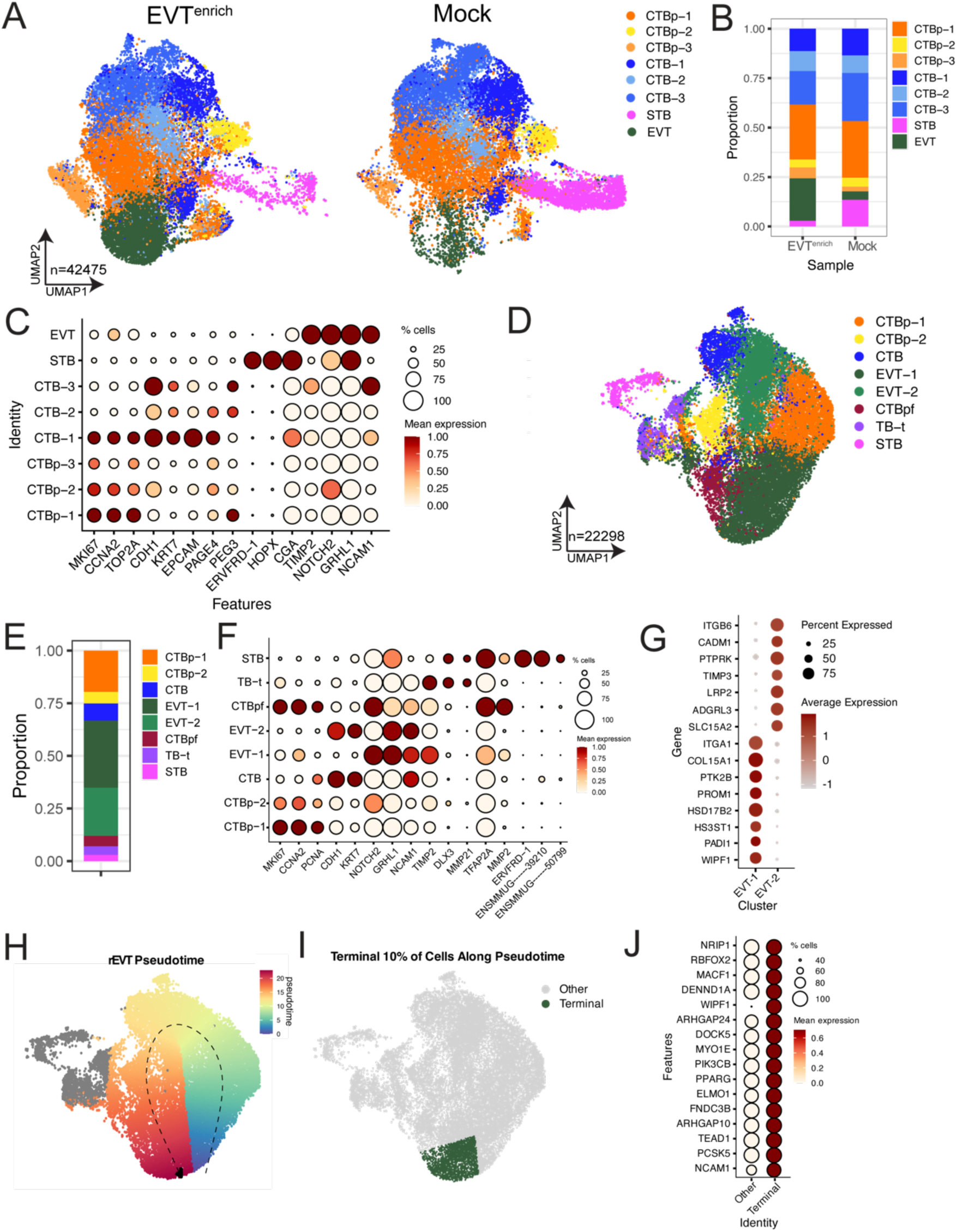
Single-nucleus RNA-seq reveals extent of EVT differentiation in rTOs. Three unique lines of rTOs induced to differentiate into EVTs (EVT^enrich^) were submitted for sn-RNAseq. **(A),** UMAP of merged datasets of EVT^enrich^ and mock samples (n= 42,475 total nuclei). **(B),** Enrichment of trophoblast cell types in integrated EVT^enrich^ and mock dataset. **(C),** Dot plot illustrating gene expression of indicated trophoblast markers. Key at right. **(D),** UMAP of cell type clusters from EVT^enrich^ rhesus trophoblast organoids (n= 22,298 nuclei). Eight clusters were identified including two proliferative cytotrophoblast clusters (CTBp-1, CTBp-2), one cytotrophoblast cluster (CTB), two extravillous trophoblast clusters (EVT-1, EVT-2), one cytotrophoblast pre-fusion cluster (CTBpf), one transitional population (tTB), and one syncytiotrophoblast cluster (STB). **(E),** Bar plot depicting enrichment of cell type clusters in EVT^enrich^ samples. **(F),** Dot plots of gene enrichment of trophoblast cell markers in EVT^enrich^ samples. **(G),** Dot plot of differentially expressed genes in EVT clusters. Key at right. **(H),** Pseudotime analysis revealed the cellular trajectory to the EVT-1 cluster. Feature plot of the pseudotime values assigned to each cell type and input into a color scale matching the UMAP based on time 0-100 EVT^enrich^ samples. **(I),** Feature plot of the terminal 10% of cells in the EVT trajectory shown in green. **(J),** Dot plot of enriched genes expressed in the terminal cells in the EVT trajectory. Key at right.

To more fully characterize trophoblast heterogeneity and differentiation trajectories under EVT-inducing conditions, snRNAseq data from the EVT^enrich^ rTOs were re-analyzed independently. UMAP projection of this subset revealed eight distinct clusters: two proliferative cytotrophoblast clusters (CTBp-1 and CTBp-2), one non-proliferative CTB cluster, a pre-fusion CTB population marked by high expression of endogenous retroviral transcripts (CTBpf), two EVT clusters (EVT-1 and EVT-2), one transitioning trophoblast population (TB-t), and a single STB cluster (**Figure 6D**). Quantification of cluster representation revealed that EVT-1 and EVT-2 together accounted for over 50% of all nuclei in the EVT^enrich^ rTO condition (**Figure 6E**), indicating successful and efficient induction of EVT differentiation across multiple rTO lines. The pre-fusion CTB cluster exhibited high expression of ERVFRD-1, consistent with a fusion-competent intermediate (**Figure 6F**). The TB-t cluster displayed intermediate expression of both CTB and EVT-associated genes, suggesting a transitional state along the EVT differentiation trajectory. Canonical marker analysis confirmed the identity of each population. Both EVT-1 and EVT-2 clusters expressed established rhesus EVT markers, including *TIMP2*, *GRHL1*, *NOTCH2*, and *NCAM1* (**Figure 6F**). *NOTCH2* and *NCAM1* were more highly expressed in EVT-1, suggesting that it represents a more mature EVT state.

To further resolve heterogeneity within the EVT lineage, differentially expressed genes were analyzed between EVT-1 and EVT-2. EVT-1 was enriched for genes associated with extracellular matrix remodeling, cell adhesion, and migration, such as *ITGA1*, *COL15A1*, *PTK2B*, and *PROM1*, suggesting a more invasive and mature phenotype (**Figure 6G**). In contrast, EVT-2 expressed higher levels of genes involved in signaling, cell polarity, and epithelial organization, including *NPAS3*, *LRP2*, *EFNA5*, and *TIMP3* (**Figure 6G**). These patterns indicate that EVT-1 and EVT-2 may represent distinct functional states or maturation stages within the EVT lineage. To determine if these transcriptional differences reflected a progression in differentiation, trajectory analysis was performed using Slingshot. The inferred lineage trajectory followed a path from CTBp-1 → EVT-2 → CTB → EVT-1, consistent with a model in which EVT-2 represents an earlier transitional state and EVT-1 a more terminal, differentiated population (**Figure 6H**). To test this, cells in the top 10% of pseudotime values along this lineage were isolated as a proxy for the most mature EVT state (**Figure 6I**). This subset was highly enriched for canonical markers of mature EVT identity, including *PCSK5*, *TEAD1*, *ELMO1*, *PPARG*, and *WIPF1* (**Figure 6J**). These results support a model in which EVT-1 corresponds to a terminal, functionally specialized state characterized by matrix remodeling and migratory potential, while EVT-2 reflects an earlier or less differentiated phase of EVT development.

### EVT Populations Are Transcriptionally Distinct Between Rhesus and Human TOs

To investigate species-specific features of EVT differentiation, we integrated snRNAseq datasets from EVT^enrich^ rTOs (rEVT) with a previously published dataset of EVT^enrich^ hTOs (hEVT)^12^. This analysis revealed eight transcriptionally distinct clusters (**Figure 7A**), with both shared and species-enriched populations. Several clusters showed a balanced representation from both species, indicating conserved transcriptional programs. CTB-1 and CTB-2 contained near-equal proportions of hEVT and rEVT cells (CTB-1: 52% hEVT / 48% rEVT; CTB-2: 57% hEVT / 43% rEVT) (**Figure 7B, S8B**). CTBp was enriched in rEVT (72%) but still contributed a substantial fraction of cells from hEVT (28%), consistent with a shared progenitor state that may differ in expansion or persistence across species. In contrast, other clusters exhibited striking species-specific enrichment. EVT-1, the dominant EVT population, was heavily skewed toward rhesus cells (69% rEVT / 31% hEVT), while EVT-2 was strongly human-enriched (95% hEVT / 5% rEVT), suggesting divergent EVT maturation pathways or regulatory programs. The STB cluster also showed species bias, with 72% of cells from hEVT and 28% from rEVT, though earlier analyses demonstrated overall conservation of STB transcriptional identity between species. Finally, CTB-4, a minor cluster, was almost exclusively derived from rEVT (93% rEVT / 7% hEVT), raising the possibility of a rhesus-specific trophoblast population or transitional state. Together, these data highlight both conserved and divergent features of trophoblast differentiation in human and rhesus organoids. While core CTB populations are largely shared, EVT subtypes show pronounced species-specific enrichment, underscoring evolutionary divergence in EVT development.

**Figure 7.**
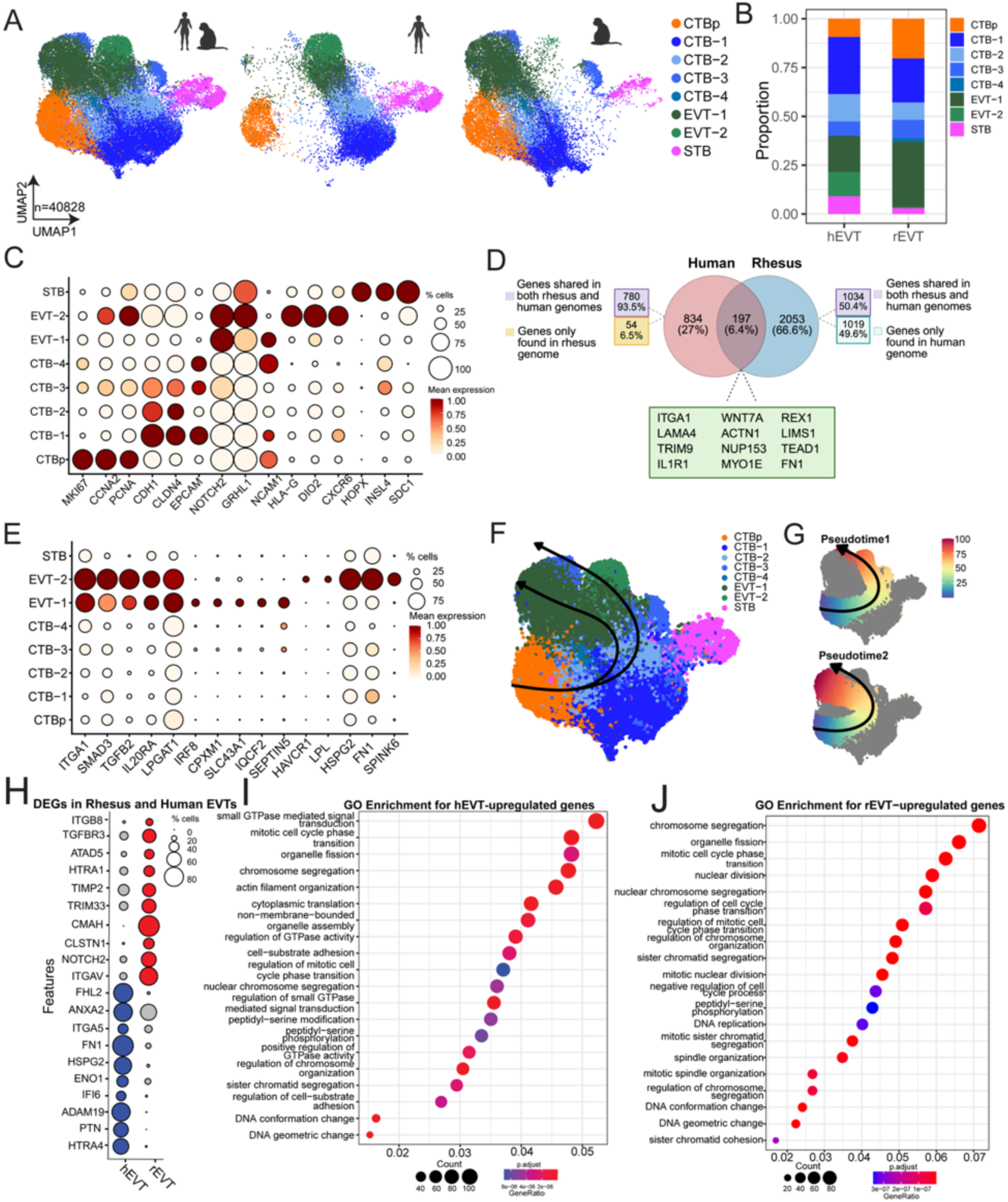
Integrated Analysis of EVT-enriched rTOs and hTOs. **(A),** UMAP of merged datasets of EVT^enrich^ rTO and EVT^enrich^ hTO datasets (n= 40,828 total nuclei). Three unique lines of EVT^enrich^ rTOs and three unique lines of EVT^enrich^ hTOs were used for analysis. Left, merged UMAP of clusters from EVT^enrich^ rTOs and EVT^enrich^ hTOs. Middle, cells from EVT^enrich^ hTOs and right, cells from EVT^enrich^ rTOs. **(B),** Enrichment of trophoblast cell types in integrated EVT^enrich^ datasets. **(C),** Dot plot illustrating gene expression of indicated trophoblast markers. Key at right. **(D),** Venn diagram of markers in the EVT^enrich^ rTO and EVT^enrich^ hTO EVT clusters that identifies shared and distinct markers. Markers that were shared are shown as genes that are present in both the rhesus macaque and human genome. Distinct rhesus and human EVT markers include genes that are shared in the rhesus macaque and human genome as well as genes that are specific to one genome. **(E),** Dot plot of enriched genes in EVT clusters in integrated dataset. **(F, G),** Pseudotime analysis of the merged of EVT^enrich^ rTO and EVT^enrich^ hTO datasets. The trajectory was mapped onto the UMAP, with pseudotime values assigned to each cell. (G) These values were then visualized using a color scale corresponding to pseudotime progression from 0 to 100 across the integrated dataset. **(H),** EVT cells were subsetted from the integrated dataset for further analysis of the EVT population. Dot plot of differentially expressed genes in rhesus and human EVTs. Key at right. **(I, J),** Dot plots of biological processes identified by differential expression between rhesus and human EVTs. GO enrichments for upregulated genes in human EVTs (I) and in rhesus EVTs (J). Key at bottom.

Analysis of cluster markers confirmed expression of canonical genes defining each trophoblast population, including CTBp markers (*MKI67*, *CCNA2*, *PCNA*), CTB markers (*CDH1*, *CLDN4*, *EPCAM*), EVT markers (*NOTCH2*, *GRHL1*, *NCAM1*, *HLA-G*, *DIO2*, *CXCR6*), and STB markers (*HOPX*, *INSL4*, *SDC1*) (**Figure 7C**). To compare EVT differentiation across species, we examined expression of EVT markers in EVT^enrich^ rTOs and hTOs. Only 6.4% of EVT markers were shared between species (**Figure 7D**), highlighting substantial divergence in EVT identity. To explore these species-specific differences, we analyzed both shared and cluster-specific markers within the EVT populations (**Figure 7E**). Shared markers included *ITGA1* and *SMAD3*, consistent with conserved roles in adhesion and TGF-β signaling. In contrast, *SLC43A1* and *SEPTIN5* were enriched specifically in EVT-1, while *HSPG2* and *FN1* were selectively expressed in EVT-2, reflecting distinct regulatory programs within each cluster. Pseudotime analysis of the integrated EVT^enrich^ rTO and hTO datasets revealed lineage trajectories to both EVT clusters (**Figure 7F**). Each trajectory originated in CTBp populations, progressed through intermediate CTB states, and terminated in its respective EVT cluster (**Figure 7G**), supporting the presence of structured differentiation routes in both species. To assess species-specific differences in EVT gene expression, we next subsetted EVT cells from the integrated dataset and performed differential expression analysis between rhesus and human EVTs (**Figure 7H**). In rhesus EVTs, *ITGB8* and *HTRA1* were significantly enriched, while human EVTs showed increased expression of *ANXA2* and *FN1*. Gene ontology enrichment analysis of these species-specific genes revealed distinct pathway associations, including the upregulation of small GTPase-mediated signal transduction in human EVTs and the upregulation of DNA conformation change in rhesus EVTs (**Figure 7I, 7J**). Together, these results underscore the molecular divergence between rhesus and human EVTs, despite shared lineage structure and induction conditions.

## Discussion

Existing *in vitro* models have failed to capture the cellular and transcriptional complexity of the rhesus macaque placenta. In this study, we established and comprehensively characterized rhesus trophoblast organoids using immunostaining, bulk RNA sequencing, and both single-cell and single-nucleus RNA sequencing, generating a detailed transcriptional map of primate trophoblast differentiation. We demonstrate that rTOs recapitulate key transcriptional signatures and secretory functions of the *in vivo* rhesus placenta. High-resolution single-cell and single-nuclei analyses revealed that rTOs give rise to distinct trophoblast lineages, enabling the identification of specific markers for rhesus CTBs, STB, and EVTs. This platform supports direct cross-species comparisons, and integration with human trophoblast organoid datasets revealed conserved STB transcriptional programs but divergent EVT identities. Pseudotime trajectory analysis further uncovered species-specific differentiation pathways along both the STB and EVT lineages. Collectively, these findings establish rTOs as a tractable model for studying primate-specific features of placentation and provide a valuable resource for investigating trophoblast biology across evolution.

Comparative analyses of select *in vitro* models of human trophoblasts and primary placental tissue have highlighted key limitations in their ability to fully recapitulate *in vivo* biology^23^. For example, while monolayer cell lines (e.g. BeWo, JEG-3, and JAR) are easy to culture, they are tumor-derived and do not spontaneously differentiate into multiple trophoblast lineages. While primary human trophoblasts do differentiate (e.g., to form fused STB), they have limited proliferation capability and a short life span in cell culture. TSC models overcome many of these limitations but have limited ability to differentiate into all trophoblast lineages and require exogenous agents for proliferative trophoblasts to differentiate into specific trophoblast populations, particularly when grown in two-dimensions. Previously described human trophoblast and decidua organoids have provided a physiologically relevant model of the maternal-fetal interface that is able to spontaneously differentiate into distinct trophoblast lineages and recapitulate the transcriptional profile of placental tissue across gestation^9,11,15^. Of note, hTOs derived from late gestation placental tissue more closely recapitulate the transcriptional signature of the first trimester placenta^11,12^. It remains uncertain whether the rTOs described here accurately replicate the rhesus macaque placenta during the first trimester. However, mCG secretion in macaques occurs only during the early stages of pregnancy and is undetectable after day 40, suggesting that rTOs most likely resemble the first trimester placenta, as we observed mCG secretion. Studies comparing rTOs to first trimester placental tissue are limited by the inherent challenges of accessing early gestation samples from nonhuman primates. Due to ethical and logistical constraints, such tissue is rarely available, and as a result, we utilized placental samples from rhesus macaques that had previously undergone experimental treatment. While these conditions were not designed specifically for placental studies, we derived multiple independent rTO lines from different animals and observed no consistent differences in organoid formation, lineage composition, or transcriptional profiles across lines. Our use of available tissue reflects the broader difficulty of conducting early pregnancy research in nonhuman primates and underscores the importance of optimizing and validating *in vitro* models like rTOs to extend the utility of these rare samples.

In both rhesus macaque and human placentas, the STB plays a critical role in nutrient exchange, hormone production, and immune protection by forming the outermost cell layer of the placenta that interfaces with maternal blood. While the overall function of the STB is conserved across both species, there are differences in structure and cellular dynamics. In humans, the STB is characterized by an extensive, highly vascularized surface area with numerous microvilli and a uniform, thin multinucleated layer, optimizing nutrient and gas exchange. In contrast, the rhesus macaque STB has a slightly less extensive surface area with a change in phenotype that occurs after implantation has begun^24^. These differences reflect species-specific adaptations in placental structure and function. Our findings demonstrate both similarities and differences in the gene expression profiles of the STB between rhesus macaques and humans. We observed the expression of canonical markers of the STB, including *HOPX* and *CGA*, in both rhesus macaque- and human-derived organoids. These markers, known for their roles in trophoblast function and placental development, were consistently present across both species, underscoring the conserved nature of STB gene expression across species. Despite conservation of select canonical markers, we also identified divergence in the transcriptional profiles of the STB between rhesus macaque and humans. For example, while *PAGE4* is a known CTB marker in the human placenta, its expression increased along the STB trajectory in rTOs^25^. Additionally, genes such as *WNT5A* and *NGF*, which are involved in STB differentiation in rhesus macaques, were not observed in humans. These genes have been shown to be upregulated in the rhesus macaque placenta compared to the human placenta^4^. These findings highlight the complex interplay between conserved functions and species-specific adaptations in the STB, which might provide valuable insights into the evolutionary pressures shaping placental biology and offering new models for further comparative studies.

Placental genes and signaling pathways are conserved along primates, but differences in gene expression have been observed associated with placental development^4^. While the placentas of rhesus macaques and humans are both hemochorial and discoid, there are differences in their development and structure. A key difference is the trophoblast invasion in the rhesus macaque placenta compared to the human placenta. EVT invasion is crucial for anchoring the placenta to the uterine wall and remodeling maternal blood vessels to ensure adequate blood supply to the fetus in both rhesus macaque and human placentas. However, in humans, EVT invasion is more extensive, penetrating deep into the decidua and myometrium, whereas in rhesus macaques, the invasion is restricted and localized, reflecting species-specific adaptations in placental structure and function^24,26^. These structural differences in EVTs may stem from distinct gene expression profiles and signaling cues between rhesus macaque and human trophoblasts. There has been no true consensus on confirmed EVT markers in the rhesus macaque placenta. Previous studies have shown that many human-associated EVT markers are present in STB cells in rhesus macaques, complicating the identification of true EVT markers in this species^14^. These findings underscore the species-specific differences in gene expression and cellular identity between rhesus macaque and human placentas. In this study, we validated *NCAM1*, *GRHL1*, and *NOTCH2* as robust EVT markers in rTOs, demonstrating that organoid models enable precise characterization of trophoblast subtypes in nonhuman primates. By leveraging these models, we identified reliable lineage markers and observed significant divergence in EVT transcriptional programs between rTOs and hTOs. While we identified strong conservation of STB identity across species, our results highlight the distinct nature of EVT differentiation pathways. By directly comparing EVT cells in rTOs and hTOs using differential expression, we observed increased expression of *ITGB8* and *HTRA1* in rhesus macaques and *ITGA5* and *FN1* in humans. The genes associated with the regulation of small GTPase mediated signal transduction pathway are also upregulated in human EVT cells and this pathway is known to play a role in EVT migration and invasion^27^. Upregulated genes in rhesus macaque EVTs were associated with the DNA geometric change pathway. Genes associated with this pathway include *HMGB1*, *HMGB3*, and *CH1D* pertinent for transcriptional changes required for EVT properties. These species-specific differences in GO enrichment for upregulated genes are consistent with the known differences in EVTs between rhesus macaques and humans. Organoid systems thus provide a critical platform for cross-species comparisons of specific trophoblast cell types, offering new insights into both conserved and divergent features of primate placental development. More broadly, these findings reflect evolutionary adaptations in trophoblast function that align with species-specific reproductive strategies and support the use of species-matched organoid models to uncover the unique regulatory mechanisms shaping placental biology.

The studies presented here establish rhesus trophoblast organoids as a tractable *in vitro* model for detailed transcriptional profiling of placental cell types. Our findings reveal cell type– specific differences between rhesus macaque and human trophoblasts, providing new insights into species-specific aspects of placental biology. The ability to directly compare human and rhesus trophoblast organoids offers a powerful platform to investigate the evolutionary adaptations that have shaped primate placentation. Leveraging these models enables dissection of both conserved and divergent mechanisms underlying placental development and function, offering a unique window into the evolution of reproductive strategies across species.

## Acknowledgements

This work was supported by the NIH grant AI145828 (C.B.C.), P01AI129859 (A.K. and C.B.C.), and the Tulane National Primate Research Center (TNPRC) NIH Base Grant P51OD011104; RRID: SCR_008167. We also gratefully acknowledge the Duke Light Microscopy Core Facility, Duke Center for Genomic and Computational Biology (GCB), Duke Proteomics and Metabolomics Core, the University of Wisconsin National Primate Research Center, and the Veterinary Medicine staff at TNPRC.

## Materials and Methods

### Rhesus macaque samples

Placental tissues were obtained at Caesarian section from seven third trimester pregnant Indian-ancestory rhesus macaques housed at the Tulane National Primate Research Center (TNPRC). The pregnant females included five rhesus cytomegalovirus (RhCMV)-seronegative dams experimentally infected with two RhCMV strains in early second trimester gestation, and two RhCMV-seropositive dams that were experimentally depleted of either CD8+ T-lymphocytes or CD20+ B-lymphocytes in late first trimester gestation. All animals remained healthy until the time of elective Caesarian section conducted between 141-154 gestation days. All macaques were from the specific pathogen free breeding colony at TNPRC and free of SIV, STLV, Type D retrovirus and herpes B virus. The study was approved by the TNPRC Institutional Animal Care and Use Committee (IACUC) and conducted in accordance with the Guide for the Care and Use of Laboratory Animals at the NIH.

### Human samples

Human tissue used in this study was obtained from Duke University after approval was received from the Duke University Institutional Review Board (IRB) and in accordance with the guidelines of Duke University human tissue procurement. Human placental tissue used in this study for the generation of organoids was collected from third trimester (38^th^-41^st^ weeks) from Caesarean sections. Tissue was collected from healthy normal donors and tissue was excluded in cases of fetal anomalies or aneuploidy.

### Derivation and culture of organoids from human and rhesus macaque placental samples

Placental tissue was carefully dissected and separated into chorionic villi, smooth chorion, amnion, and decidua separated. The amnion layer was kept intact and peeled from the chorion. To remove decidua from the chorion, isolated chorion tissue was carefully scraped with forceps. Stem/progenitor cells isolation from smooth chorion were similar to full-term chorionic villous isolation previously described^11^. Briefly, dissected tissue was cut into small pieces (1-2mm^2^) and sequentially digested with 0.2% trypsin-250 (Alfa Aesar, J63993-09)/0.02% EDTA (Sigma-Aldrich E9884-100G) and 1.0 mg/mL collagenase V (Sigma-Aldrich, C9263-100MG) on a 37°C stir plate in small bottles containing stir bars. After the collagenase V digestion, tissue was disrupted manually by vigorous pipetting with a 5-10mL serological pipette (10-20 times) prior to filtration. Then pool two digests flow-through and pellet by centrifugation (600g for 6 minutes) and finally resuspend with ice-cold growth-factor-reduced Matrigel (Corning 356231). Matrigel “domes” (40ul/well) were plated into 24-well tissue culture plates (Corning 3526), placed in a 37°C incubator to pre-polymerize for approximately 3 minutes, turned upside down to ensure equal distribution of the isolated cells in domes for another 10 minutes, then carefully overlaid with 500 µL prewarmed media. For both rTOs and hTOs derivation and culture, the term trophoblast organoid medium (tTOM) consisting of Advanced DMEM/F12 (Life Technologies, 12634-010) supplemented with 1× B27 (Life Technologies, 17504-044), 1× N2 (Life Technologies, 17502-048), 10% FBS (vol/vol, Cytiva HyClone, SH30070.03), 2 mM GlutaMAX^TM^ supplement (Life Technologies, 35050-061), 100 µg/mL Primocin (InvivoGen, ant-pm-1), 1.25 mM N-Acetyl-L-cysteine (Sigma, A9165), 500 nM A83-01 (Tocris, 2939), 1.5 µM CHIR99021 (Tocris, 4423), 50 ng/mL recombinant human EGF (Gibco, PHG0314), 80 ng/mL recombinant human R-spondin 1 (R & D systems, 4645-RS-100), 100 ng/mL recombinant human FGF2 (Peprotech, 100-18C), 50 ng/mL recombinant human HGF (Peprotech, 100-39), 10mM nicotinamide (Sigma, N0636-100G), 5 µM Y-27632 (Sigma, Y0503-1MG), and 2.5 µM prostaglandin E2 (PGE2, R & D systems, 22-961-0) was used. To passage, TOs were digested in prewarmed TrypLE Express (Gibco, 12605-028) at 37°C for 8 min. Further mechanical dissociation was achieved by manual disruption. Dissociated TOs were collected and washed by centrifuge, then resuspended in fresh ice-cold Matrigel and replated as domes at the desired density for continuous culture. For freezing, overlaid media was aspirated, and organoid-Matrigel domes were resuspended in CryoStor CS10 stem cell freezing medium (STEMCell Technologies, 07930) and frozen at -80 °C and then transferred to liquid nitrogen for long-term storage. For thawing cryopreserved TOs, organoids were thawed as quickly as possible, diluted with 5 times volume of basic TOM containing Advanced DMEM/F12, 2 mM GlutaMAX supplement, 10 mM HEPES (Gibco, 15630-106), 1 × Penicillin/Streptomycin (Lonza, 17-602E) and centrifuged to pellet. Afterwards, TOs were resuspended in new ice-cold Matrigel and replated for recovery culture and passaged as described above.

### Derivation and culture of organoids from human and rhesus macaque decidua samples

Decidual tissue was dissected from human and rhesus macaque placenta and further cleaned by removing as much as possible those attached chorionic villi prior to the isolation of decidua glands-enriched suspension as described previously (Turco et al., 2017b; Turco et al., 2017a, Yang et al., 2022). Briefly, collected decidual tissues were cut into small pieces then digested in 1.25 U/mL Dispase II (Sigma-Aldrich, D4693)/0.4mg/mL collagenase V (Sigma-Aldrich, C-9263). Following digestion and pipetting, the decidua glandular structures were filtered out with 100 μm cell strainer, then collect the glandular structures from strainer and pipette to dissociate them into small pieces before resuspending with ice-cold growth-factor-reduced Matrigel (Corning 356231), plated as “domes” in 24-well plates (Corning, 3526), and overlaid with 500 µL prewarmed media. For decidua organoids Expansion Medium (ExM) was used with the same composition as previously described^28^. ExM was renewed every 2-3 days. Mature decidua organoids were passaged by mechanical disruption following TrypLE Express (ThermoFisher Scientific, 12605010) digestion.

### Generation of conditioned media

Overlaid media was collected from mature organoids after 7-10 days post-passaging. CM was stored at -20°C. Non-conditioned medium (NCM) was TOM (described above) that had not been exposed to organoids. All experiments using CM were from at least two independent TO lines.

### Hormone Secretion Assays

Conditioned media was collected as described above and a radioimmunoassay was used to quantify mCG and progesterone secretion in cell conditioned media as previously described^18,19^.

### Mass Spectrometry

CM was collected from rTOs and submitted to Duke University Proteomics and Metabolomics Core Facility. SDS-PAGE gel bands were subjected to in-gel tryptic digestion using standardized protocols^29^. Digested peptides were lyophilized to dryness and resuspended in 12 uL of 0.2% formic acid/2% acetonitrile. Each sample was subjected to chromatographic separation on a Thermo Neo Vanquish UPLC equipped with a 1.7 µm HSS T3 C18 75 µm I.D. X 250 mm reversed-phase column (NanoFlow data). The mobile phase consisted of (A) 0.1% formic acid in water and (B) 0.1% formic acid in acetonitrile. 3 µL was injected and peptides were trapped for 3 min on a 5 µm Symmetry C18 180 µm I.D. X 20 mm column at 5 µl/min in 99.9% A. The analytical column was then switched in-line and a linear elution gradient of 5% B to 40% B was performed over 30 min at 400 nL/min. Data collection on the Fusion Lumos mass spectrometer with a FAIMS Pro device was performed for three difference compensation voltages (−40v, −60v, −80v). Within each CV, a data-dependent acquisition (DDA) mode of acquisition with a r=120,000 (@ m/z 200) full MS scan from m/z 375 - 1500 with a target AGC value of 4e5 ions was performed. MS/MS scans with HCD settings of 30% were acquired in the linear ion trap in “rapid” mode with a target AGC value of 1e4 and max fill time of 35 ms. The total cycle time for each CV was 0.66s, with total cycle times of 2 sec between like full MS scans. A 20s dynamic exclusion was employed to increase depth of coverage.

Raw LC-MS/MS data files were processed in Proteome Discoverer 3.0 (Thermo Scientific) and then submitted to independent Sequest database searches against a Rhesus Macaque protein database containing both forward (20260 entries) and reverse entries of each protein. Search tolerances were 2 ppm for precursor ions and 0.8 Da for product ions using trypsin specificity with up to two missed cleavages. All searched spectra were imported into Scaffold (v5.3, Proteome Software) and scoring thresholds were set to achieve a peptide false discovery rate of 1% using the PeptideProphet algorithm. Raw data were also imported into Protein Discoverer 3.0 for area under the curve measurements of eluting peptides.

### Immunofluorescence

Organoids were fixed in 4% PFA for 15 min at room temperature, then washed in PBS. Organoids were placed in 0.25% Triton X-100/PBS to permeabilize for 15 min at room temperature. Organoids were washed in 1x PBS and then incubated with the indicated primary antibodies in PBS at 4°C overnight. Organoids were pelleted by gravity, washed with PBS, and then incubated for 30 min at room temperature with Alexa Fluor-conjugated secondary antibodies (Invitrogen). Organoids were washed again with PBS and mounted in Vectashield (Vector Laboratories) containing 4′,6-diamidino-2-phenylindole (DAPI) and were transferred onto microscope slides with large-orifice 200 µL-tips (Fisher Scientific, 02707134). The following antibodies and reagents were used: cytokeratin-19 (Abcam, ab9221, ab52625), SDC-1 (abcam, ab128936), ITGA6 (Invitrogen, MA5-16884), E-cadherin (Invitrogen, PA5-85088), Mucin 5AC (Abcam, ab3649), Alexa Fluor 594–conjugated phalloidin (Invitrogen, A12381).

### Induced Differentiation of rTOs into EVT^enrich^ rTOs

To generate EVT-enriched rTOs, we adapted established EVT differentiation protocols developed previously^11,13,22^. rTOs were passaged as described above and plated in Matrigel domes into 24-well tissue culture plates. Organoids were cultured in TOM as described above for 3-4 days, then switched to EVT differentiation media 1 [EVT m1: advanced DMEM/F12 supplemented with 2 mM L-glutamine, 0.1 mM 2-mercaptoethenol, 0.5% (vol/vol) penicillin/streptomycin, 0.3% (vol/vol) BSA, 1% (vol/vol) ITS-X supplement, 100 ng/ml NRG1 (Cell Signaling, 5218SC), 7.5 µM A83-01, 2.5 µM Y27632 and 4% (vol/vol) Knockout Serum Replacement (Gibco 10828010)], for 6 days. EVT m1 was renewed every 2 days for the initial 4 days then daily for the remaining culture period. Following, organoids were switched to EVT m2, which is the same recipe as described above for EVT m1 but without NRG1, for a further 2-3 days, with daily media changes.

### RNA extraction and Bulk RNA-seq

Total RNA was isolated with the Sigma GenElute total mammalian RNA miniprep kit or Qiagen RNAeasy Mini Kit following the manufacturer’s instruction and using supplementary Sigma DNase digestion. RNA quality and concentration were determined using a Nanodrop ND-1000 Spectrophotometer. For bulk RNA-seq analysis, RNA was isolated from organoids or placenta tissue as described above. Purified Total RNA was verified by Thermo scientific Nanodrop one. The libraries were prepared by the Duke Center for Genomic and Computational Biology (GCB) using the Tru-Seq stranded total RNA prep kit (Illumina). Sequencing was performed on the NovaSeq 6000 by using 150-bp paired-end sequencing. The reads were aligned to the human or rhesus macaque reference genome (GRCh38, Mmul10) using QIAGEN CLC Genomics (v20). Differential expression analysis was performed using the DeSeq2 package in R ^30^, k-means pathway cluster enrichment for GO Biological processes and principal component analyses (PCA) was performed using iDEP ^31,32^. Heatmaps and hierarchical clustering was performed in R using the pheatmap package and were based on log_2_ RPKM (reads per kilobase million). Files associated with bulk RNA-seq studies have been deposited into Sequence Read Archive (PRJNA891648 and PRJNA1230745). Principal component analyses were performed using pcaExplorer in R ^33^. Volcano plots were generated using the EnhancedVolcano package in R or in Graphpad Prism version 9.

### Dissociation of organoids for single cell and single nucleus RNA sequencing

Organoids were dissociated by scraping Matrigel domes into 1 mL of pre-warmed TrypLE Express (Invitrogen, 12605036) and incubating at 37°C for 12 min, swirling the tube every 2-3 min. Dissociated organoids were pelleted at 1250 rpm for 3 min and re-suspended in 200 µL DMEM containing 10% FBS. Resuspended organoids were subjected to vigorous manual disruption using a single channel p200 pipette (Ranin, 17008652) for 3 min followed by the addition of 800 µL of DMEM containing 10% FBS. The disrupted suspension was then passed over a 40 µm filter cell strainer (Corning, 352098). Flow through was then centrifuged at 1250 rpm for 5min and the pellet resuspended in 250 µL of 1x PBS for a final volume of ∼300 µL and cell counts of ∼1 x 10^6^ cells/mL. Cell viability was determined using trypan blue and followed by automated cell counting (Biorad TC20 automated cell counter). Sequencing was performed on three organoid lines from unique placental tissue.

### Single-Cell RNA-seq Library Preparation and Data Analysis. SC Processing

Libraries for single-cell RNA sequencing were prepared from 10k total cells using the methods described in the 10x Genomics Single Cell 3’ Reagent Kit Protocol (v2 chemistry, Manual Part #CG00052). Single-cell RNA sequencing runs were performed on an Illumina NovaSeq 6000 (Illumina, San Diego) using an S2 flow cell (Illumina, San Diego), which allowed for ∼61k reads/cell for rTO samples. hTO sequencing was performed using an S4 flow cell (Illumina, San Diego), which allowed for ∼74k reads/cell.

### SN Processing

Nuclei were isolated from dissociated rTOs using 10x Genomics’ Nuclei Isolation from Single Cell Suspensions protocol (10x Genomics – Pleasanton, VA). All buffers used were prepared fresh, according to manufacturer’s protocol. Cell suspensions were pelleted by centrifugation at 300rcf, 4°C for 5 minutes, followed by removal of the supernatant, and lysis buffer was added with gentle pipetting on ice for 5 minutes. Next, samples were washed twice with nuclei wash and resuspension buffer, followed by centrifugation at 500 rcf, 4°C for 10 minutes then supernatant removal. Nuclei are then resuspended in wash and resuspension buffer, concentatrion determend on a Cellometer Ascend Automated Cell Counter (RevvityWaltham, MA). If necessary, nulei are filtered through 40µm Flowmi Cell Strainers (Bel-Art – Wayne, NJ) to remove clumps and debris prior to proceeding with the single nuclei assay. Nuclei are visualized at 40x magnification using LifeTech EVOS FL microscope (Thermo Fisher Scientific – Waltham, MA) to determine quality via nuclear membrane intactness. Nuclei were loaded with a Chromium Reagent Kit v4. Libraries were sequenced on the Nova Seq X Plus at a targeted depth of 22,000 to 35,000 reads per nucleus.

### SC and SN Sequencing Data Processing

Post-processing, quality control, and read alignment to the Mmul10 rhesus macaque reference genome or to the hg38 human reference genome were performed using 10x CellRanger package (v6.1.2, 10x Genomics). Gene expression matrices generated by the 10x CellRanger aggregate option were analyzed using Seurat (version 4.0) in R ^34–37^. For SC, cells with at least 200 and no more than 10,000 unique expressed genes were included in downstream analysis, and cells with more than 10% mitochondrial reads were excluded from analysis. Dropout rate was 67-74%. For SN, nuclei with at least 200 and no more than 8,500 unique genes and less than 50,000 counts were included in downstream analysis, and nuclei with more than 1% mitochondrial reads and 5% ribosomal reads were excluded from analysis. Dropout rate was 87-89%. To eliminate batch effects, datasets from unique donors were normalized using the *sctransform* function and integrated using the *FindIntegrationAnchors()* in Seurat (version 4.0) in R ^37^. Dimensional reduction was performed using the *RunPCA()* function to obtain the first 20 principal components, which was determined using *ElbowPlots()* across the first 50 dimensions. To identify clusters, Louvain clustering (Seurat *FindClusters()* function) was performed, and optimal resolution was determined using the *clustree()* function^38^ on a range of resolutions between 0.2-1.0, with 0.3-0.4 used for downstream analyses. Integration of rTO and hTO datasets was performed on previously integrated datasets using the Harmony algorithm^39^ (v1.0) for batch correction and alignment. Differential expression analysis between clusters was performed using the Wilcoxon rank sum test implemented in Seurat’s FindAllMarkers() function. Genes with a log2 fold change > 0.25 and an FDR-adjusted p-value < 0.05 were considered significantly differentially expressed. Marker genes for each cluster were identified by searching for positively enriched genes using the aforementioned criteria. All files associated with this single cell and single nuclei RNA-seq study have been deposited into Sequence Read Archive (PRJNA1230239 and PRJNA1283913).

The *slingshot* package (version 2.6.0) in R was used to determine differentiation trajectories from clusters identified in Seurat with unbiased starting and ending roots^40^. The raw counts and above-generated slingshot object were used to run *evaluateK()* with the total number of knots ranging from 3 to 9. The optimal number of knots was determined to be 4. The *fitGAM()* function using *tradeSeq* (1.5.10) was run with this resulting value and gene expression along lineages identified using the *associationTest()* function^41^. Heatmaps of expression changes across lineages were generated using *ComplexHeatmap()* on log transformed counts and rasterized using the *ImageMagick* “Bessel” filter ^42^. The *plotGenePseudotime()* function was used to visualize raw count gene expression in individual cells across lineages from the *slingshot* object. Code used for data analysis and figure generation are available at CoyneLabDuke Github repository (https://github.com/CoyneLabDuke/Rhesus-Macaque-Trophoblast-Organoids).

### Statistics and reproducibility

All experiments reported in this study have been reproduced using independent samples (tissues and organoids) from multiple donors. Data are presented as mean ± SD, unless otherwise stated. Statistical significance was determined as described in the figure legends. Parametric tests were applied when data were distributed normally based on D’Agostino-Pearson analyses; otherwise, nonparametric tests were applied. For all statistical tests, p value <0.05 was considered statistically significant, with specific p values noted in the figure legends.

## Supplemental Figure Legends

**Supplemental Figure 1.**
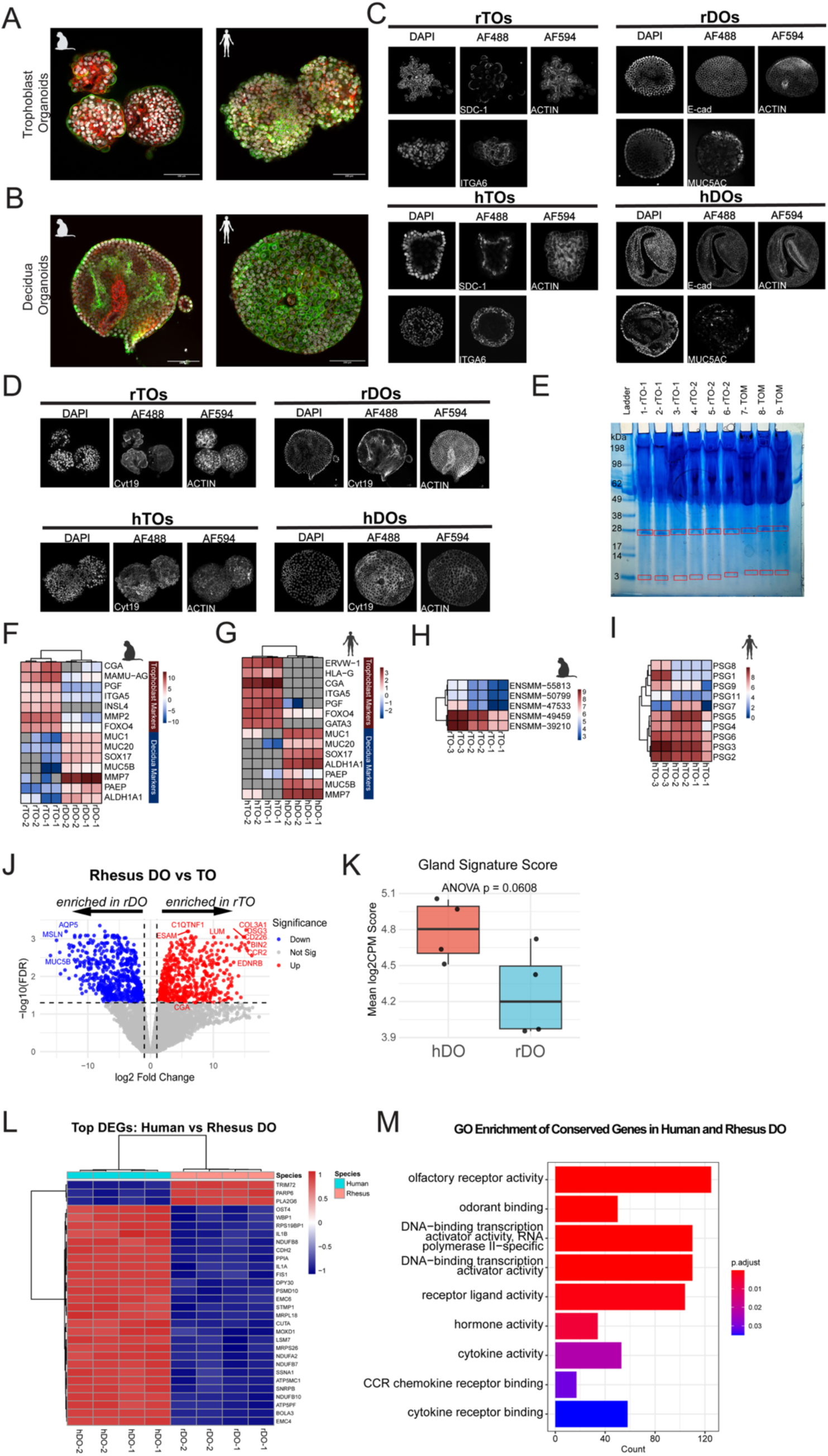
Validation of rTOs and rDOs. **(A, B),** Confocal micrographs of rTOs (A) and hTOs (B) immunostained for CYT-19 (green), ACTIN (red), DAPI-stained nuclei are in black and white. Scale bar, 100μm. **(C),** Confocal micrographs of immunostained rTOs, hTOs, rDOs and hDOs in black and white. **(D),** Confocal micrographs of rDOs and hDOs immunostained for CYT-19 (green), ACTIN (red), DAPI-stained nuclei are in black and white. **(E),** SDS-PAGE gel of rTO CM samples. Red boxes indicate bands that were excised and used for mass spectrometry. **(F, G),** Heatmap (based on log_2_ RPKM values) of transcripts expressed in hDOs and hTOs (F) and rDOs and rTOs (G) that are associated with markers of trophoblasts and decidua. Data shown is 2 biological replicates and 2 technical replicates in rTOs and rDOs. **(H),** Heatmap (based on log_2_ RPKM values) of identified pregnancy-specific glycoproteins (PSGs) transcripts expressed in rTOs. Data shown is 3 biological replicates and 2 technical replicates in rTOs. **(I),** Heatmap (based on log_2_ RPKM values) of known pregnancy-specific glycoproteins (PSGs) transcripts expressed in hTOs. Key at right. Red indicates high levels of expression; blue indicates low levels of expression. Data shown is 3 biological replicates and 2 technical replicates in hTOs. **(J),** Volcano plot of differentially expressed transcripts in rTOs and rDOs. Red circles represent transcripts significantly upregulated and blue circles represent transcripts significantly downregulated (log_2_fold-change > ± 1 and FDR < 0.05) and gray circles represent transcripts that were not significantly changed. **(K),** Boxplot of gland signature score in hDOs and rDOs determined by glandular epithelial markers. **(L),** Heatmap of top differentially expressed genes in hDOs and rDOs. Key at right, red indicates high levels of expression and blue indicates low levels of expression. **(M),** Bar plot of GO enrichment terms associated with conserved genes in hDOs and rDOs. Key at right.

**Supplemental Figure 2.**
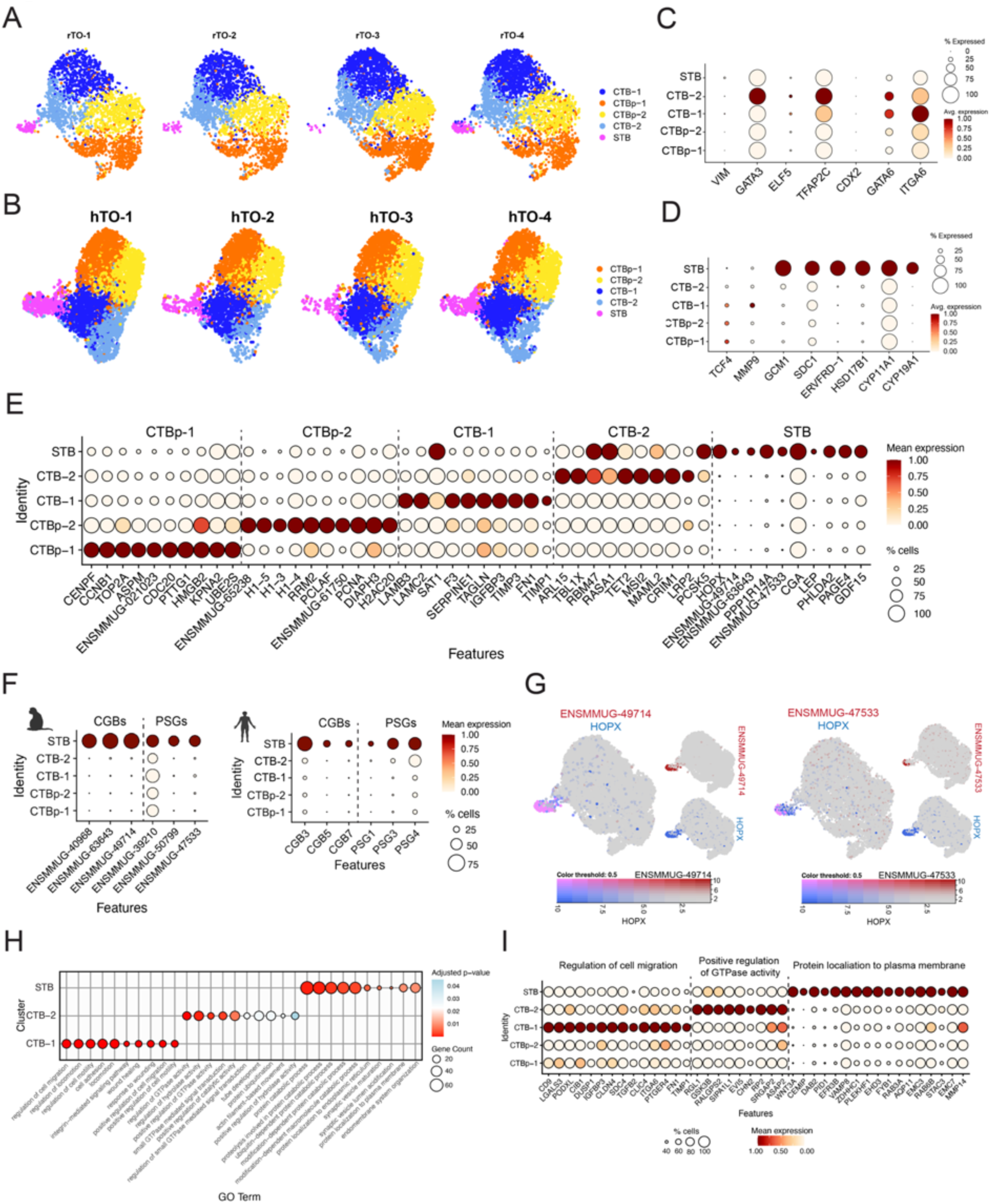
Defining cell types in rTO and hTO sc-RNAseq datasets. Individual UMAPs from samples of **(A)** Four rTO samples and **(B)** Four hTO samples. Dot plots of gene expression of **(C)** cytotrophoblast markers and **(D)** syncytiotrophoblast or extravillous trophoblast markers in rTOs. **(E),** Dot plot of top 10 markers in each cluster in rTOs. Key at right. **(F),** Dot plot of rhesus CGBs and PSGs in rTOs (left) and human CGBs and PSGs in hTOs (right). Key at right. **(G),** Combined feature plots of rhesus CGB (ENSMMUG00000049714) and HOPX (left) and rhesus PSG (ENSMMUG00000047533) and HOPX (right) in rTOs. Key at bottom. **(H),** Top biological processes for CTB-1, CTB-2, and STB clusters in rTOs identified using ClusterProfiler. **(I),** Dot plots of genes associated with top biological processes in CTB-1, CTB-2 and STB clusters in rTOs.

**Supplemental Figure 3.**
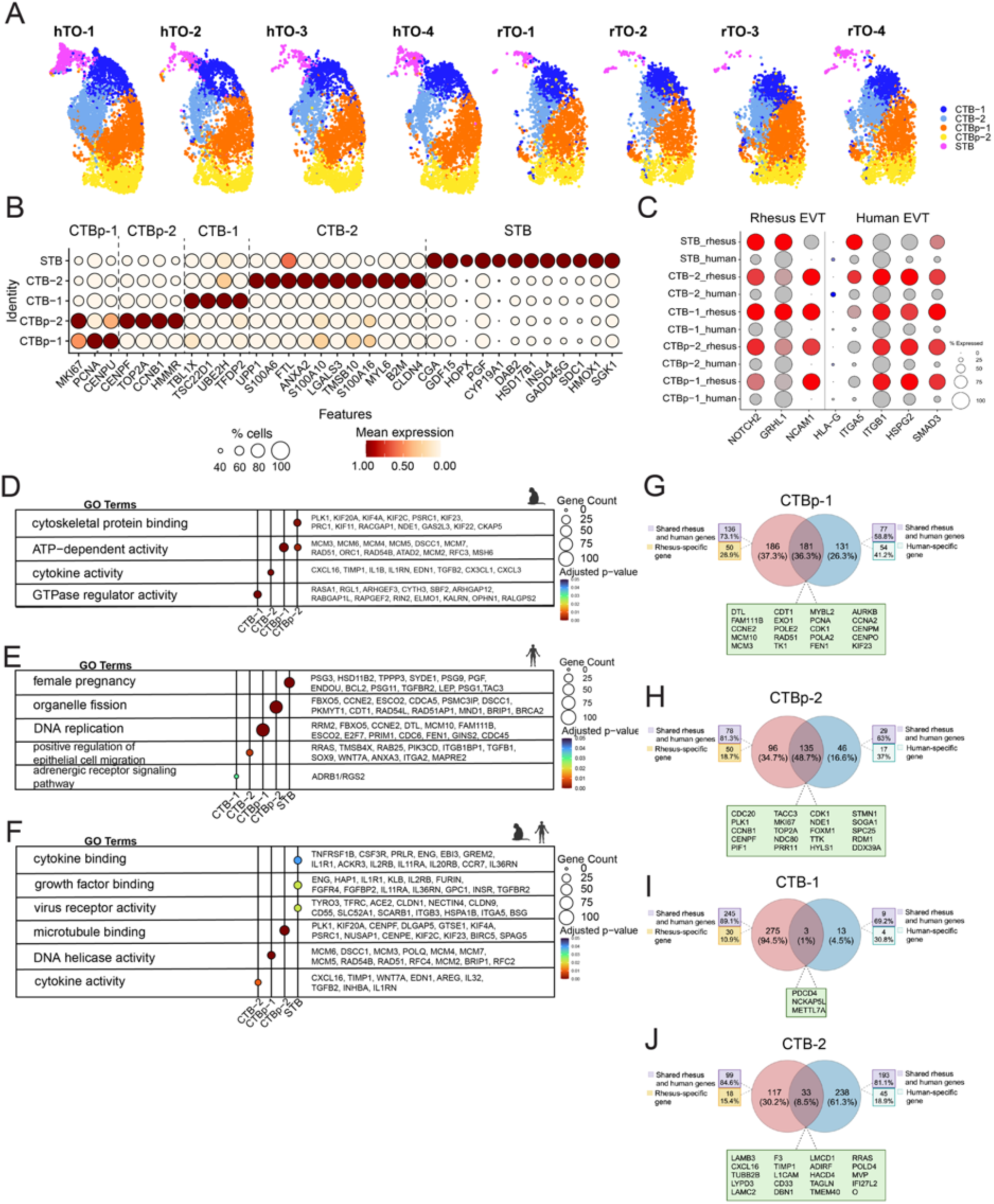
Defining cell types in integrated sc-RNAseq dataset with rTO and hTO. **(A),** Individual UMAP of hTO and rTO samples that comprise that integrated dataset. **(B),** Dot plot of gene expression of cytotrophoblasts, proliferative cytotrophoblasts, and syncytiotrophoblasts. **(C),** Dot plot of rhesus macaque and human extravillous trophoblast markers in the integrated dataset (rhesus are shown in red and human shown in blue). **(D, E, F),** Subset of pathways from GO term enrichment in all clusters in subsetted (D) rTO dataset and (E) hTO (F) integrated rTO and hTO dataset. Venn diagram of markers in the rhesus and human **(G),** CTBp-1, **(H),** CTBp-2, **(I),** CTB-1, **(J),** CTB-2 clusters that identifies shared and distinct markers. Markers that were not shared are shown as genes that are present in both the rhesus macaque and human genome as well as markers that are specific to one genome.

**Supplemental Figure 4.**
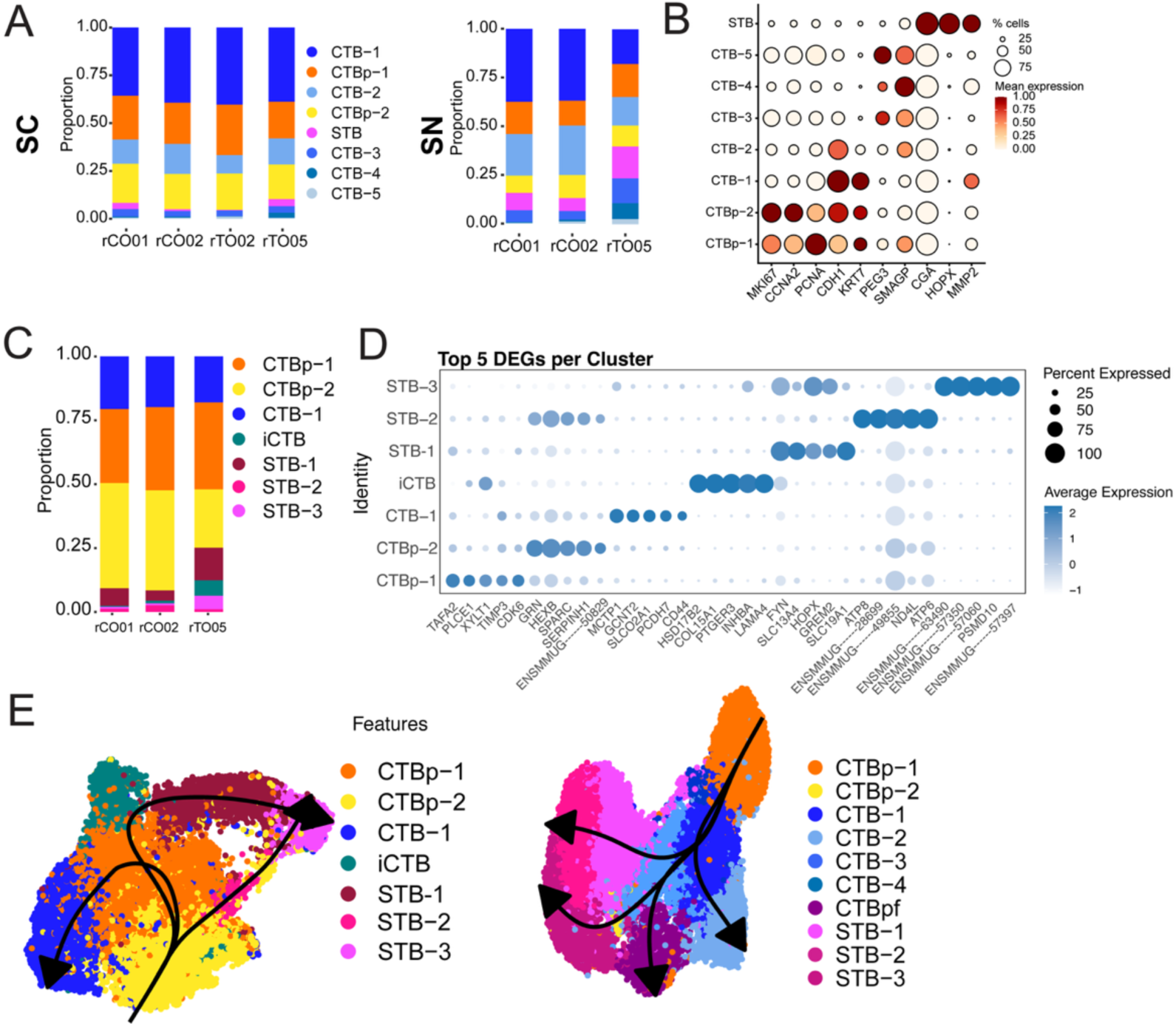
Defining cell types in rTO sn-RNAseq and integrated sn-RNAseq and sc-RNAseq datasets. **(A),** Individual bar plots showing enrichment of cell types in sc-RNAseq (SC) samples (left) and sn-RNAseq (SN) samples (right). **(B),** Dot plot of expression of canonical trophoblast markers in integrated SC and SN dataset. **(C),** Cell type enrichment of individual rTO samples. **(D),** Dot plot of top five differentially expressed genes per cluster in rTO sn-RNAseq dataset. Key at right. **(E),** Pseudotime analysis of the rTO (F) and hTO (G) datasets. The trajectories were mapped onto their respective UMAPs.

**Supplemental Figure 5.**
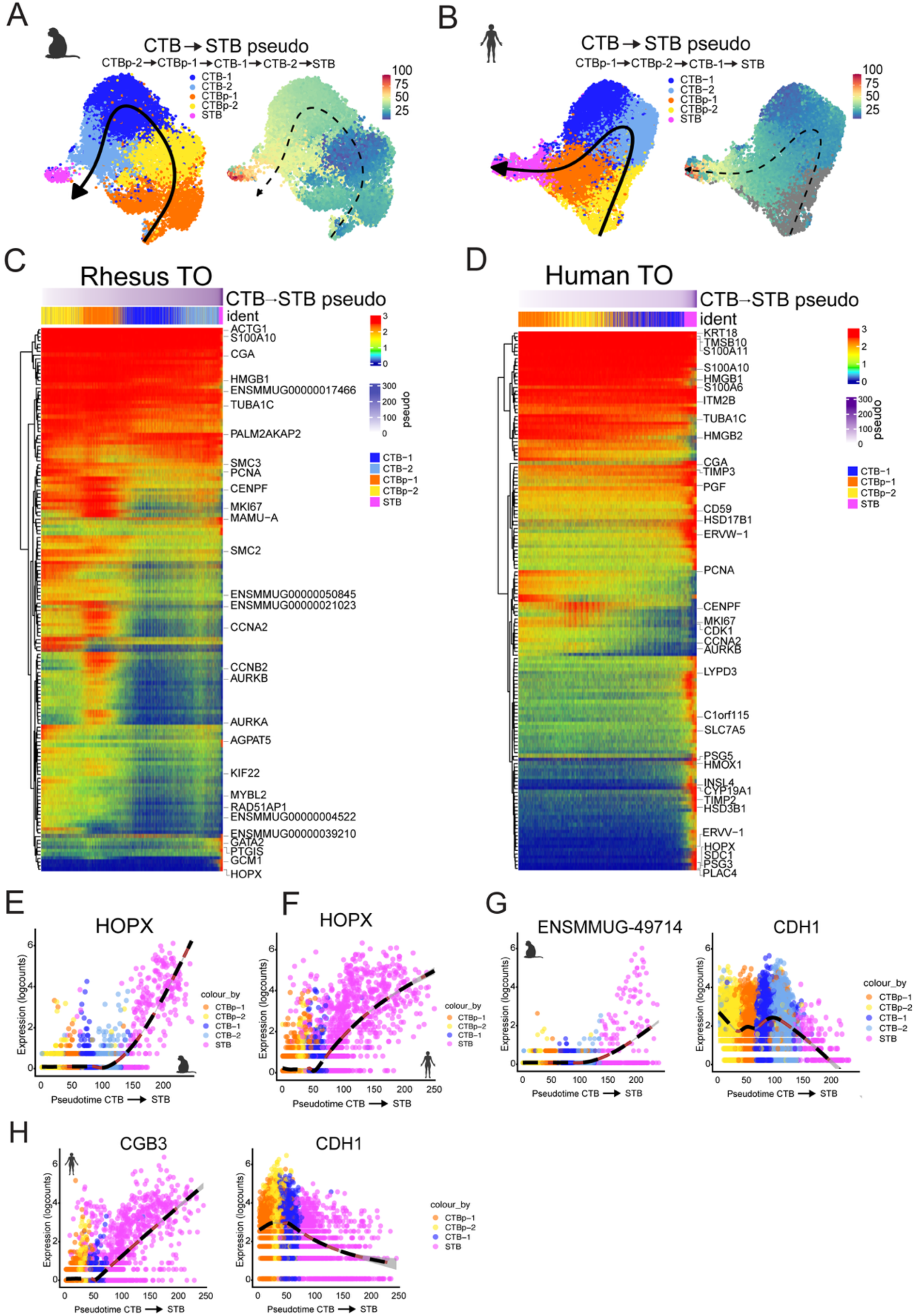
Syncytiotrophoblast cell differentiation trajectory in rhesus and human trophoblast organoids. **(A,B),** Pseudotime was performed using Slingshot analysis to define cellular lineage reconstruction across sc-RNAseq datasets. This illustrates the trajectory of the cell types in the clusters in rTO (A, right) and hTO (B, right). Pseudotime values were assigned to each cell type and input into a color scale matching the UMAP based on time 0-100 in rTO (A, left) and hTO (B, left). The arrow across the UMAP shows the order of the trajectory. **(C, D),** Pseudotime values revealed top genes correlated to the trajectory in rTO (C) and hTO (D). **(E, F),** HOPX gene expression increases along the differentiation trajectory to STB subtype in rTO (E) and hTO (F). The cell types presented are represented by the colors denoted in the legend. Logcounts and pseudotime values are shown as a color scale present in the legend. **(G),** Normalized expression of ENSMMUG-49714and CDH1 shown along rTO pseudotime. **(H),** Normalized expression of CGB3 and CDH1 shown along hTO pseudotime.

**Supplemental Figure 6.**
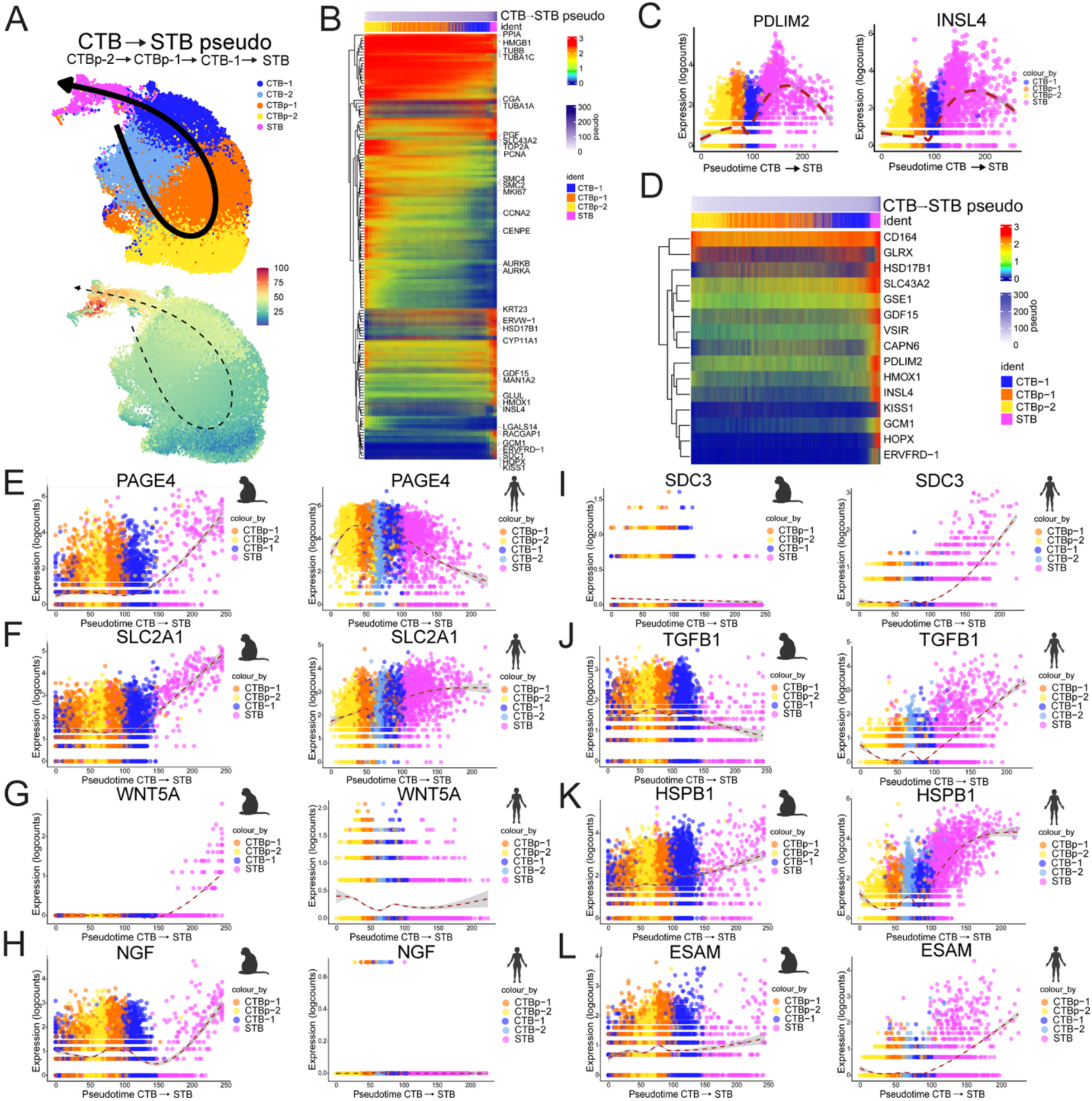
Comparative single cell RNA-seq reveals distinct STB differentiation trajectories between human and rhesus macaque trophoblast organoids. **(A),** Pseudotime analysis of the integrated rTO and hTO datasets. The trajectory was mapped onto the UMAP, with pseudotime values assigned to each cell. These values were then visualized using a color scale corresponding to pseudotime progression from 0 to 100 across the integrated dataset. **(B),** Heatmap of the top genes associated with the pseudotime enriched in the combined rTO and hTO trajectory dataset. **(C),** Normalized expression of PDLIM2 and INSL4 shown along integrated pseudotime in the combined dataset. **(D),** Heatmap of genes that increase along the trajectory to the STB subtype in the integrated dataset. The cell types presented are represented by the colors denoted in the legend. Logcounts and pseudotime values are shown as a color scale present in the legend. **(E, F, G, H),** Normalized expression of PAGE4, SLC2A1, WNT5A, and NGF shown across rTO (on left) and hTO (on right) trajectories. **(I, J, K, L),** Normalized expression of SDC3, TGFB1, HSPB1, and ESAM shown along rTO (on left) and hTO (on right) trajectories.

**Supplemental Figure 7.**
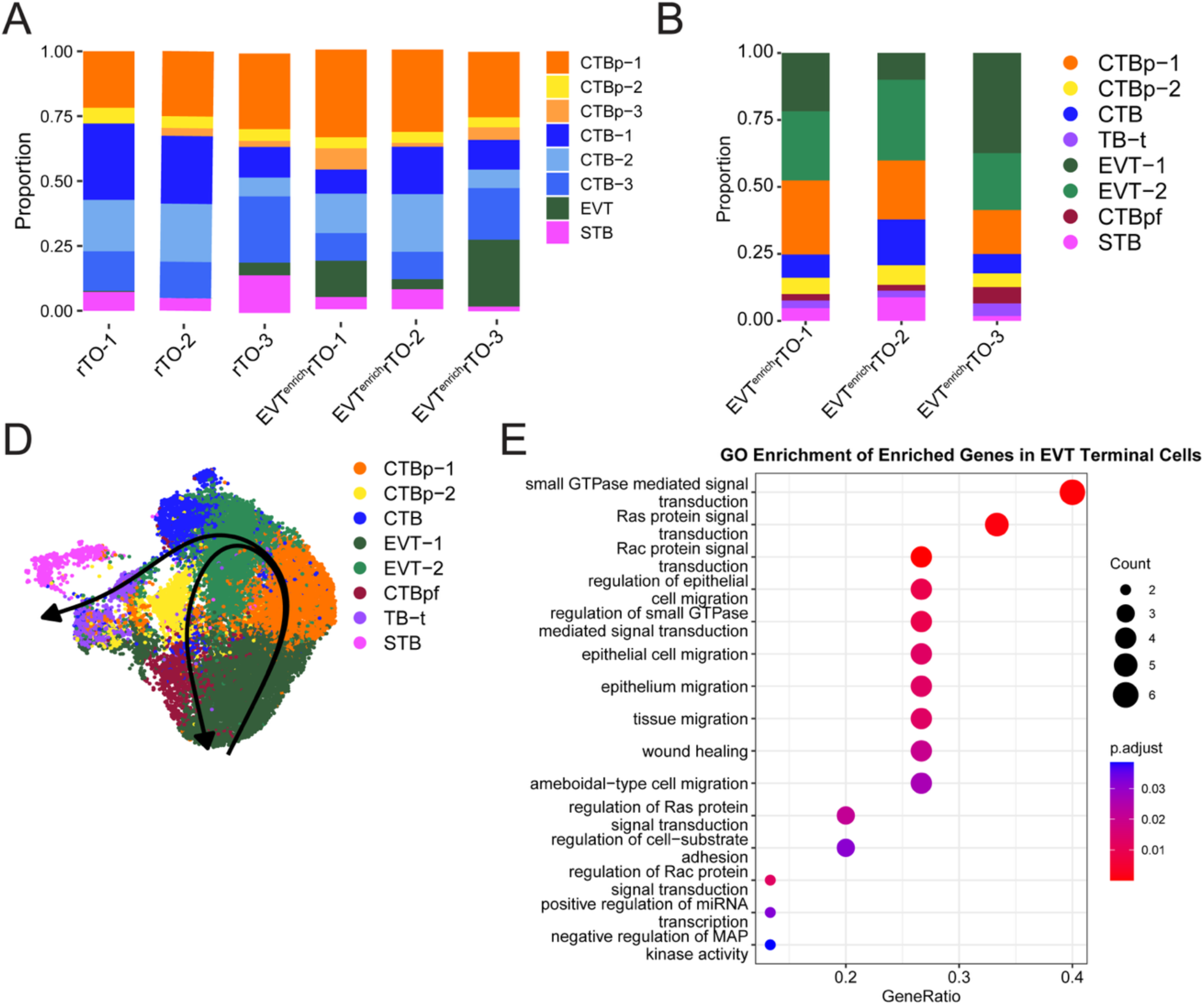
Defining Cellular Identities in Rhesus EVT^enrich^ datasets. **(A),** Bar plots of cluster enrichment from integrated dataset of Mock and EVT^enrich^ samples. **(B),** Bar plots of cluster enrichment in EVT^enrich^ samples. **(C),** Pseudotime analysis of the EVT^enrich^ dataset. The trajectories were mapped onto the UMAP. **(D),** Dot plot of GO enrichment of enriched genes in 10% terminal cells in EVT trajectory compared to other cells.

**Supplemental Figure 8.**
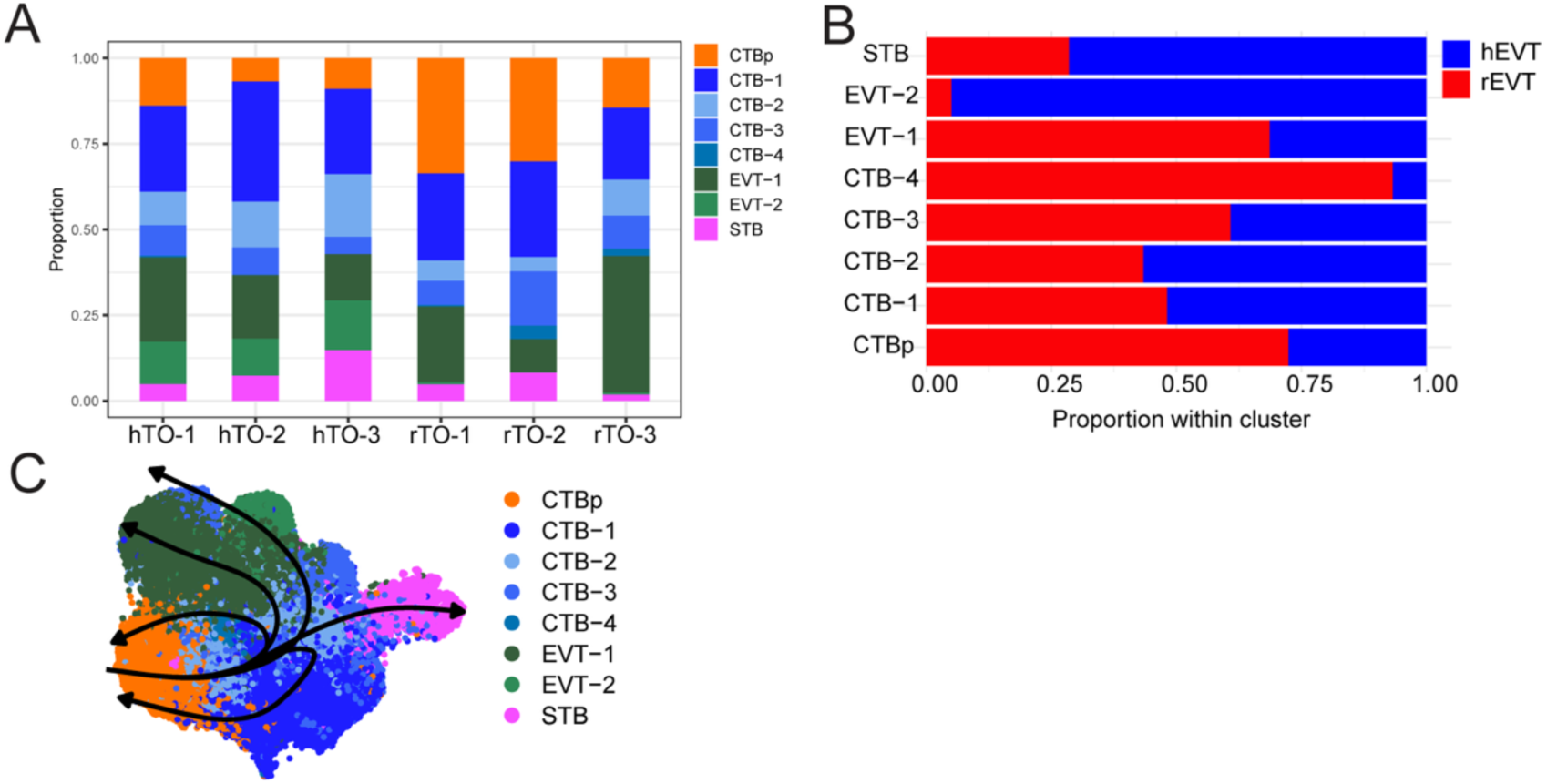
Defining Trophoblast Population in Integrated Human and Rhesus EVT^enrich^. **(A),** Cluster enrichment of individual samples in merged dataset. **(B),** Bar plot of proportion of cluster comprised of human (in blue) or rhesus nuclei (in red). **(C),** Pseudotime analysis of the integrated EVT^enrich^ dataset. The five trajectories were mapped onto the UMAP.

## Supplemental Table Legends

**Supplemental Table 1.** Metadata of animal study design.

| Tissue Code |  |
| --- | --- |
| 01 | CMV-seropositive healthy pregnancy;<br>CD8b depleted at 55 days gestation,<br>~141 days gestation age- elective C-<br>section |
| 02 | CMV-seronegative dams inoculated with<br>FL-RhCMV + RhCMV UCD52 in early<br>2<sup>nd</sup> trimester and C-section near term<br>gestation |
| 03 |  |
| 04 |  |
| 05 |  |
| 06 |  |
| 07 | CMV-seropositive healthy pregnancy;<br>Anti-CD20 antibody administered for B<br>Lymphocyte depletion at 53 days<br>gestation and elective C-section at 144<br>days gestation |

**Supplemental Table 2.** Metadata of trophoblast organoid lines.

| Experiment | TO code | Gestational age | TOs sex | TO passage number |
| --- | --- | --- | --- | --- |
| Bulk RNA-seq of rTOs | rTO02 | 154d | F | P3 |
|  | rCO02 | 154d | F | P3 |
|  | rCO01 | 141d | F | P2 |
| Bulk RNA-seq of rDOs | rDO01 | 141d | F | P5 |
|  | rDO02 | 154d | F | P2 |
| Bulk RNA-seq of hTOs | TO74 | 41w | M | P5 |
|  | CO80 | 40w | F | P3 |
|  | CO82 | 40w | M | P3 |
| Bulk RNA-seq of hDOs | DO77 | 40w | F | P4 |
|  | DO76 | 40w | F | P5 |
| Mass Spectrometry | rCO02 | 154d | F | P3, P4, P7 |
|  | rCO05 | 143d | F | P3, P6, P7 |
| Radioimmunoassay | rTO02 | 154d | F | P3, P5, P6, P7 |
|  | rTO05 | 143d | F | P6, P7 |
|  | rTO04 | 149d | F | P4 |
| scRNAseq of rTOs | rCO01 | 141d | F | P3 |
|  | rCO02 | 154d | F | P2 |
|  | rTO02 | 154d | F | P4 |
|  | rTO05 | 143d | F | P3 |
| scRNAseq of hTOs | CO80 | 40w | F | P5 |
|  | TO68 | 27w | M | P12 |
|  | CO82 | 40w | M | P4 |
|  | CO81 | 40w | F | P4 |
| snRNAseq of rTOs | rCO01 | 141d | F | P5 |
|  | rCO02 | 154d | F | P12 |
|  | rTO05 | 143d | F | P4 |
| snRNAseq of EVT-differentiated rTOs | rCO01 | 141d | F | P2 |
|  | rCO02 | 154d | F | P3 |
|  | rTO05 | 143d | F | P7 |

**Supplemental Table 3.**
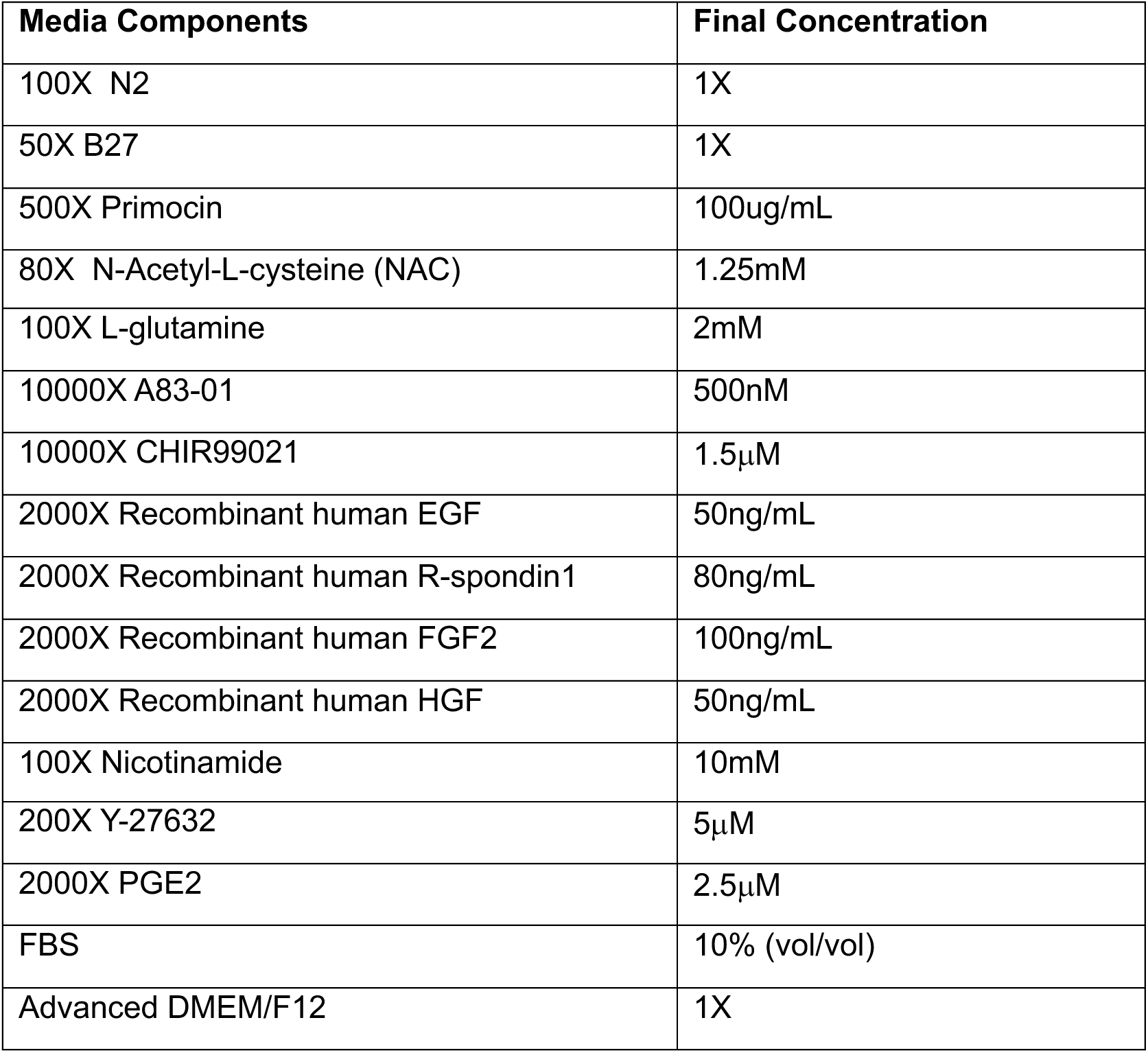
Components of term trophoblast organoid (tTOM) media.

**Supplemental Table 4.**
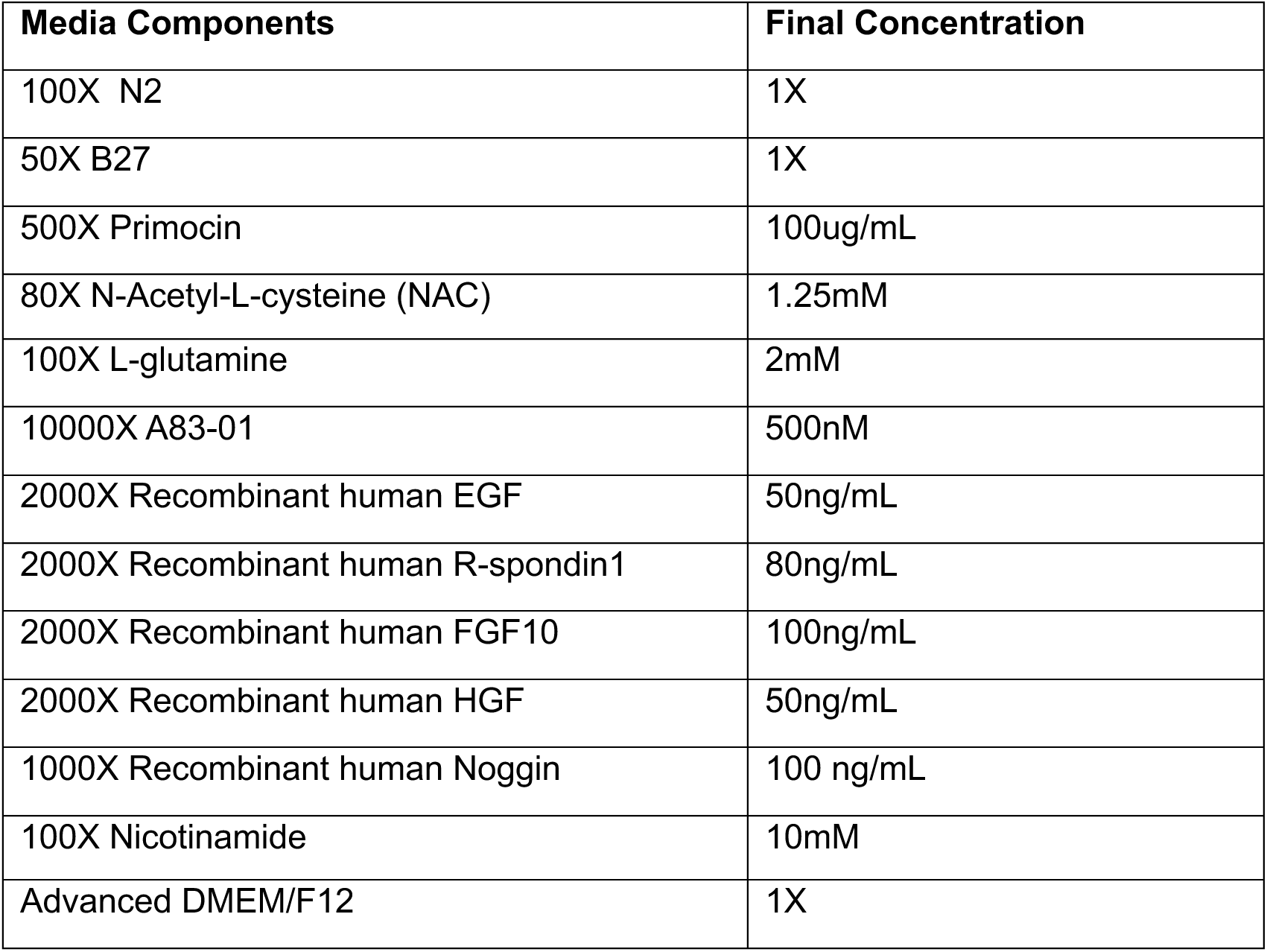
Components of decidua organoid media.

**Supplemental Table 5.**
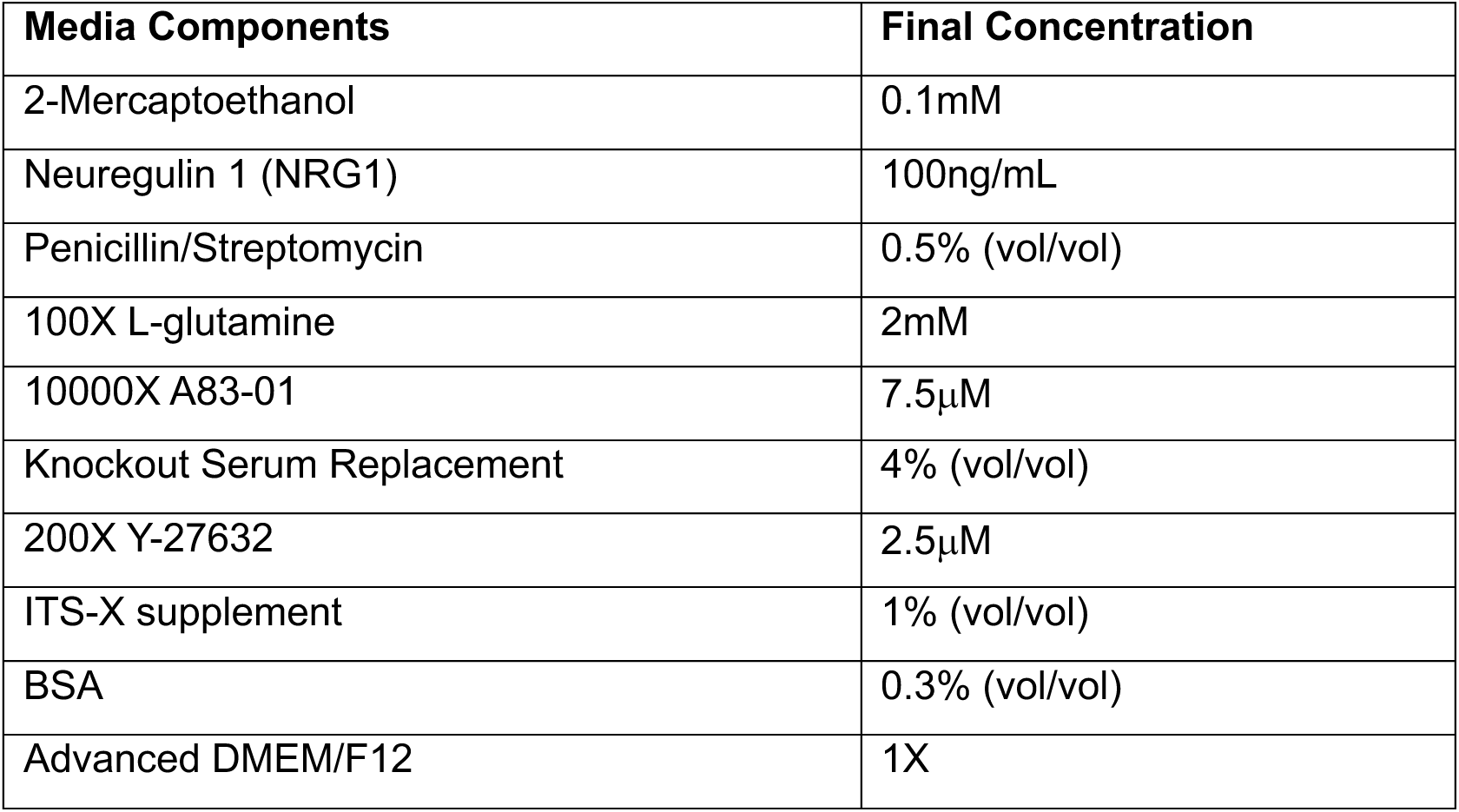
Components of EVT differentiation media 1.

**Supplemental 6.**
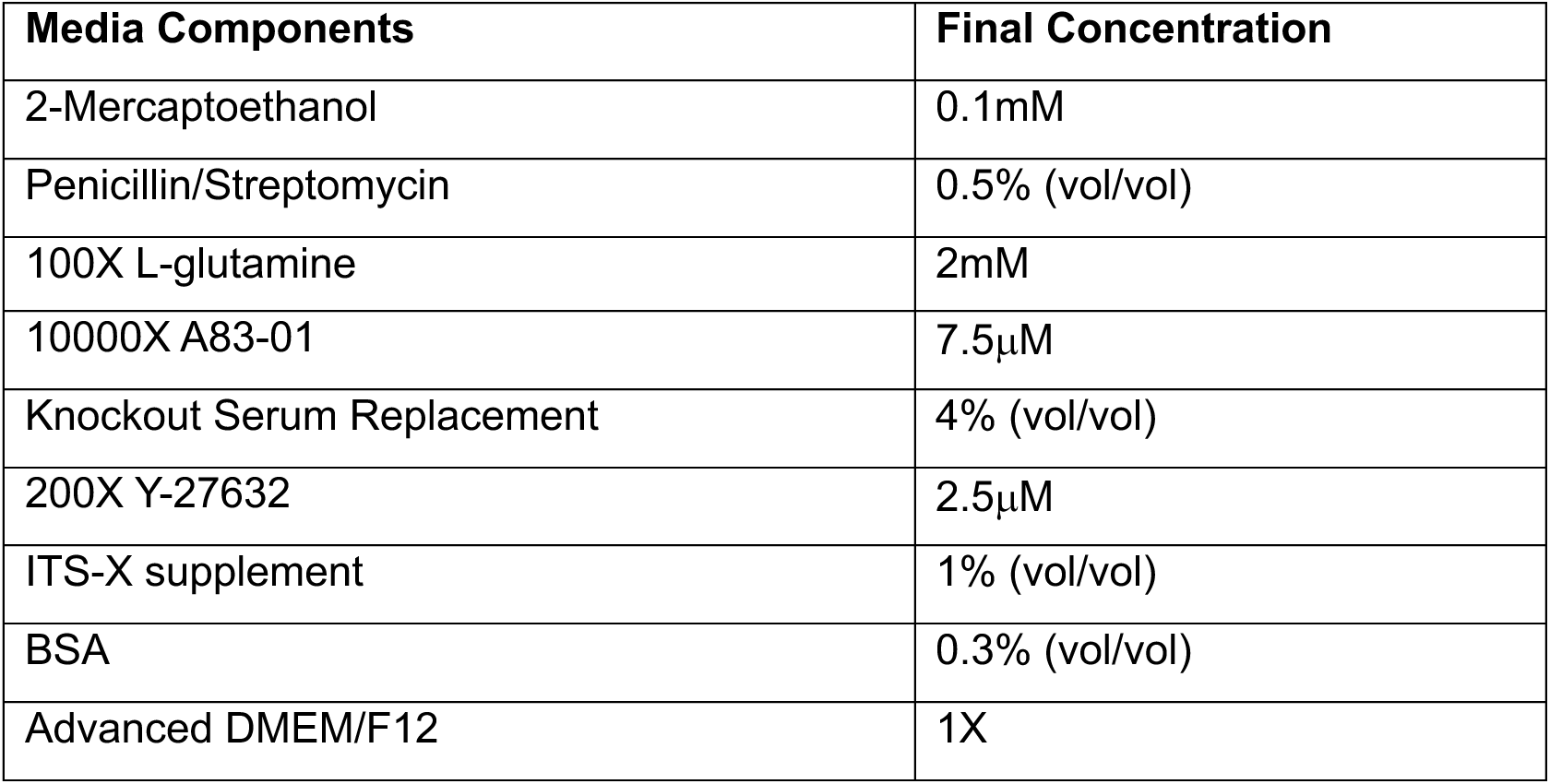
Components of EVT differentiation media 2.

**Supplemental Table 7.** Antibodies used for immunofluorescence staining.

| Antibody | Source | Catalog number | Additional Information |
| --- | --- | --- | --- |
| Mouse monoclonal Cytokeratin-19 | Abcam | ab9221 | 1:500 |
| Rabbit monoclonal Cytokeratin-19 | Abcam | ab52625 | 1:500 |
| Mouse monoclonal Mucin 5AC | Abcam | ab3649 | 1:500 |
| Rabbit polyclonal E-cadherin | Invitrogen | PA5-85088 | 1:500 |
| Rabbit monoclonal SDC-1 | Abcam | ab128936 | 1:500 |
| ITGA6 |  |  |  |
| Alexa Fluor 594-conjugated phalloidin | Invitrogen | A12381 | 1:1000 |
| Alexa Fluor 488 Goat Anti-mouse IgG secondary antibody | Invitrogen | A32723 | Secondary antibody 1:1000 |
| Alexa Fluor 594 Goat Anti-mouse IgG secondary antibody | Invitrogen | A11032 | Secondary antibody 1:1000 |
| Alexa Fluor 594 Goat Anti-rabbit IgG secondary antibody | Invitrogen | A11037 | Secondary antibody 1:1000 |

**Supplemental Table 8. Significant Genes along STB trajectory in rTO snRNAseq dataset.** Genes with significant association to STB lineage trajectory identified using *associationTest()* in R. False Discovery rate (FDR) and adjusted p-value <0.05 were used to determined significant genes. These genes reflect the transcriptional changes throughout the developmental trajectory.

**Supplemental Table 9. Significant Genes along STB trajectory in rTO scRNAseq dataset.** Genes with significant association to STB lineage trajectory identified using *associationTest()* in R. False Discovery rate (FDR) and adjusted p-value <0.05 were used to determined significant genes. These genes reflect the transcriptional changes throughout the developmental trajectory.

**Supplemental Table 10. Significant Genes along STB trajectory in hTO snRNAseq dataset.** Genes with significant association to STB lineage trajectory identified using *associationTest()* in R. False Discovery rate (FDR) and adjusted p-value <0.05 were used to determined significant genes. These genes reflect the transcriptional changes throughout the developmental trajectory.

**Supplemental Table 11. Significant Genes along STB trajectory in hTO scRNAseq dataset.** Genes with significant association to STB lineage trajectory identified using *associationTest()* in R. False Discovery rate (FDR) and adjusted p-value <0.05 were used to determined significant genes. These genes reflect the transcriptional changes throughout the developmental trajectory.

## Supplemental Movie Legends

**Supplemental Movie 1. Three-dimensional reconstruction of rhesus trophoblast organoid.** Rhesus trophoblast organoids were fixed and immunostained for SDC-1 (green), a marker of syncytiotrophoblasts, and actin (red) to visualize cellular architecture. Nuclei were counterstained with DAPI (blue). Z-stack images were acquired by confocal microscopy and reconstructed into a three-dimensional rendering using Imaris software. The movie illustrates the complex organization and surface morphology of the organoids in three dimensions.

**Supplemental Movie 2. Three-dimensional reconstruction of rhesus decidua gland organoid.** Rhesus decidua gland organoids were fixed and immunostained for E-cadherin (green) and actin (red) to visualize cellular architecture. Nuclei were counterstained with DAPI (blue). Z-stack images were acquired by confocal microscopy and reconstructed into a three-dimensional rendering using Imaris software. The movie illustrates the complex organization and surface morphology of the organoids in three dimensions.

